# Spatio-temporal dynamics of microglia phenotype in human and murine cSVD: impact of acute and chronic hypertensive states

**DOI:** 10.1101/2023.08.15.553403

**Authors:** Lorena Morton, Philipp Arndt, Alejandra P. Garza, Solveig Henneicke, Hendrik Mattern, Marilyn Gonzalez, Alexander Dityatev, Deniz Yilmazer-Hanke, Stefanie Schreiber, Ildiko R. Dunay

## Abstract

Vascular risk factors such as chronic hypertension are well established major modifiable factors for the development of cerebral small vessel disease (cSVD). In the present study, our focus was the investigation of cSVD-related phenotypic changes in microglia in human disease and in the spontaneously hypertensive stroke-prone rat (SHRSP) model of cSVD. Our examination of cortical microglia in human post-mortem cSVD cortical tissue revealed distinct morphological microglial features specific to cSVD. We identified enlarged somata, an increase in the territory occupied by thickened microglial processes, and an expansion in the number of vascular-associated microglia. In parallel, we characterized microglia in a rodent model of hypertensive cSVD along different durations of arterial hypertension, i.e., early chronic and late chronic hypertension. Microglial somata were already enlarged in early hypertension, whereas at late-stage chronic hypertension they further exhibited elongated branches, thickened processes, and a reduced ramification index, mirroring the findings in human cSVD. An unbiased multidimensional flow cytometric analysis revealed phenotypic heterogeneity among microglia cells within the hippocampus and cortex. At early-stage hypertension, hippocampal microglia exhibited upregulated CD11b/c, P2Y12R, CD200R, and CD86 surface markers. Detailed analysis of cell subpopulations revealed a unique microglial subset expressing CD11b/c, CD163, and CD86 exclusively in early hypertension. Notably, even at early-stage hypertension, microglia displayed a higher association with cerebral blood vessels. We identified several profound clusters of microglia expressing distinct marker profiles at late chronic hypertensive states. We further detected a temporal hypertension-related disturbances in blood-brain barrier integrity, accompanied by increased recruitment of leukocytes to the brain parenchyma in early hypertension. In summary, our findings demonstrate a higher vulnerability of the hippocampus, stage-specific microglial signatures based on morphological features, and cell surface protein expression in response to chronic arterial hypertension. These results indicate the diversity within microglia sub-populations and implicate the subtle involvement of microglia in cSVD pathogenesis.

## Introduction

Chronic arterial hypertension, especially in midlife, has been associated with an increased risk for all-cause dementia, comprising the neurodegenerative Alzheimer’s disease (AD) and vascular cognitive impairment (VCI). Arterial hypertension causes sporadic hypertensive cerebral small vessel disease (cSVD), which in turn is responsible for the development of VCI, and for the majority of extensive intracerebral hemorrhages [35, 54, 56, 72]. In the clinic, small vessel disease is currently diagnosed through downstream, mostly irreversible, brain pathologies such as lacunes, white matter hyperintensities (WMH), or hemorrhages [73], which are considered secondary to initial microvascular dysfunction. The absence of specific biomarkers and suitable translational imaging measures hinders the bedside detection of early dysfunction. In contrast, initial cSVD-induced microvascular dysfunction has been related to chronic low-grade oxidative microvascular injury with consequent structural and functional adaptations such as blood-brain barrier (BBB) disruption and neuroinflammation [37, 53, 68]. Corroborative evidence from autopsy and molecular imaging studies supports the emergence of neuroinflammation as a major contributor to the pathogenesis of hypertensive cSVD [14, 64, 70, 71, 77]. Since neuroinflammation also plays a major role in AD pathogenesis, this phenomenon may be a critical link between neurovascular and neurodegenerative diseases [74].

Microglia are the primary cellular players in neuroinflammation [21, 26, 61]. These brain-resident immune cells respond to peripheral low-grade inflammation through excessive glial activation by increasing microvascular permeability, hence disrupting the BBB, as it has been shown in murine models developing autoimmune diseases and systemic inflammation [4, 22]. This transduction of inflammation, characterized by pro-inflammatory cytokines and chemokines commonly found in AD and VCI including cSVD, can also induce adhesion molecule expression on brain endothelial cells, leading to leukocyte infiltration, synaptic pruning, and demyelination [10, 13, 34]. Notably, in hypertensive cSVD, regions of increased microglial reactivity have been observed, which do not coincide with areas of heightened BBB permeability, suggesting a spatial dissociation between these two processes [70]. Microglia are highly dynamic and shape neuronal responses through purinergic mechanisms by interacting with both neurons and blood vessels. Altered microglial activity and related inflammatory changes can impair cerebral blood flow, and changes in neurovascular coupling were shown to precede symptom onset in several neurodegenerative disorders. Recent evidence suggests that microglial response to capillary lesions is dependent on the P2Y12 purinergic receptor (P2Y12R) for BBB repair as it was observed in studies, where inhibition of these receptors decreased microglial process motility [36, 49]. Furthermore, P2Y12R accumulation at endothelial cell contacts is essential for vasodilation, and focal loss of P2Y12R has been associated with synaptic degeneration, emphasizing the link between vascular inflammation and synaptic/neural dysfunction [12]. Still, the mechanisms of how microglia interact with the vasculature and react during peripheral low-grade chronic inflammation in hypertensive cSVD is not yet elucidated.

Thus, the spatial, temporal, and causal relationship between microglial reactivity, BBB leakages, and peripheral low-grade inflammation in hypertensive cSVD remains unanswered. Although microglial positron emission tomography studies detected a correlation between BBB disruption and increased neuroinflammation in AD and cSVD [70], the causal relationship between these events is not fully understood. Research about microglial dynamics in hypertensive cSVD must address the question, whether low-grade peripheral inflammation triggers central neuroinflammation or vice versa or if both are mutually reinforcing. Furthermore, recent evidence has to be taken into account that suggest dynamic changes in the diversity of microglia throughout life, with a high heterogeneity during early development, a more homogenous population in adulthood, and re-emergence of heterogeneity during senescence and under pathological conditions. For example, such microglia diversity could be detected in an animal model for AD, in which microglial clusters with distinct gene expressional profiles were identified in cortical regions [31, 41, 55, 66].

We here investigated phenotypical microglial features in the non-transgenic Spontaneously Hypertensive Stroke-Prone Rat (SHRSP), a rodent model of hypertensive cSVD, and in human post-mortem brain tissue of elderly with cSVD. The SHRSP rodent model develops chronic hypertension, mimicking the multimorbidity, and potentially polygenetic susceptibility, of hypertensive patients prone to cSVD [2, 3, 25, 58], as well as chronic sterile low-grade inflammation attributed to elevated arterial hypertension [15]. Here, we therefore studied morphological features of microglia and quantified vascular-associated microglia (VAMs) in both the SHRSP model of cSVD and human cSVD. To understand the role of microglia in the pathophysiology and progression of cSVD, we further examined the BBB, characterized microglia in early- and late-stage hypertension and correlated morphologic al microglial features with hypertension-induced surface marker expression profiles in the SHRSP rodent model of cSVD.

## Materials and Methods

### Animals

All experiments were performed in accordance with the German Animal Welfare Ordinance (as of August 2013) for scientific laboratory animals. Animal experiments were performed with the approval of the Animal Care Committee of Saxony-Anhalt (license identification number 42502-2-1561 Uni MD). A total of 32 male SHRSP (Charles River Laboratories, Wilmington, Massachusetts, USA) aged 25 and 34 weeks and 32 male age-matched Wistar rats (Charles River Laboratories, Research Models and Services, Germany GmbH, Sulzfeld, Germany) as controls were included in the study. From this point forward, Wistar rats are designated as controls, and SHRSP rats as early (25 weeks) and late (34 weeks) chronic hypertensive rats. All animals were housed with a natural light-night cycle and had access to water and food *ad libitum*. SHRSP rats develop a vascular risk profile characterized by arterial hypertension between 6-8 weeks of age [27, 29, 50, 58]. The animals’ health status was monitored by a daily assessment of their neurological functions (such as decreased spontaneous activity, coordination failure, and hunched posture) and supported by weekly body weight control. All animals were neurologically unremarkable during the observation period.

### Human subjects

Brain tissue of 9 human autopsy cases were included into the study. Brains of all subjects underwent routine neuropathological examination and were screened for tauopathies, alpha-synucleinopathies and beta-amyloid (Aβ) deposition. Inclusion criteria for cSVD cases for entering the study were pathologically confirmed white matter lesions (WML). Control cases were randomly selected within the same age range as the cSVD cases. Exclusion criteria for all cases were a history of neurodegenerative disorders; i.e., Alzheimer-related tau pathology exceeding Braak’s neurofibrillary stage II [5] or other tauopathy, Parkinson’s disease except incidental subcortical Lewy bodies in the lower brain stem (stage <3) according to Braak et al [7], multisystem atrophy or other alpha-synuclein-related pathology. Altogether 4 cases with cSVD (3 females, 1 male) and 5 control cases (3 females, 2 males) were investigated. The age range of the two groups presented as Mean ± SD were 64.0 ± 16.0 in the cSVD group and 63.7 ± 12.7 in the control group. Further demographics and relevant data for the patient cohort are provided in **Suppl. Table 1**. This retrospective study was performed in compliance with the University Ethics Committee guidelines as well as German federal and state law governing human tissue usage. Informed written permission was obtained from all patients and/or their next of kin for autopsy.

### Immunofluorescence in rodent tissue

Animals were transcardially perfused under pentobarbital anesthesia (40 mg/kg body weight i.p.) with 120 mL of phosphate buffered saline followed by perfusion with 120 mL of the fixative 4 % paraformaldehyde (PFA) within 8 min. Brains were removed, immersion-fixed in 4 % PFA for 48 h, cryoprotected in 30 % sucrose for 6 days, and frozen in methyl butane at −80°C. Using a cryostat, brains were sectioned from the frontal to the occipital pole, and 30 µm-thick free-floating sections were collected in PBS. Immunofluorescence staining was performed as previously described [58]. Two coronal brain sections were stained, containing the hippocampus and retrosplenial cortex of both hemispheres per animal (coronal sections Bregma -2.5 to -4.5 [19]). Sections were repeatedly washed in phosphate-buffered saline (PBS), blocked with 10 % donkey serum/0.5 % Triton X-100 (Sigma, St Louis, MO, USA), and incubated overnight at 4 °C with STL-FITC (dilution 1:750, solanum tuberosum lectin-fluorescein isothiocyanate, endothelial marker, Vector Laboratories FL-1161) and goat anti-IBA1 (dilution 1:1500, ionized calcium-binding adapter molecule 1, microglial marker, Novus Biologicals NB100-1028). Sections were incubated for two hours the following day with donkey anti-goat Cy5-conjugated secondary antibodies (dilution 1:500, Jackson ImmunoResearch, 703-175-155) and mounted on slides with Fluoromount Aqueos Mounting Medium (Merck, F4680).

### Histology and neuropathological evaluation in human post-mortem tissue

Routine neuropathological investigations were performed using hematoxylin & acid fuchsine (modified H&E), advanced silver stains, and single- and double-label immunohistochemistry as previously described [16, 75]. Briefly, brains were fixed in a 4 % solution of formaldehyde and cut in approximately 1 cm-thick coronal slabs. Tissue slabs and blocks of the frontal, mid-hemispheric, and occipital regions, the cerebellum and various brainstem regions (rostral medulla, pontine-mesencephalic junction and midbrain) were embedded in polyethylene glycol (PEG 1000, Merck, Carl Roth Ltd, Karlsruhe, Germany). Multiple 100 µm thick consecutive sections were obtained from each embedded tissue block with the aid of a sliding microtome (Jung, Heidelberg, Germany). Enlargement of subcortical perivascular spaces, hyalinosis of subcortical perforating vessels in the white matter and basal ganglia area as well as presence of WML were assessed in the modified H&E stain [16]. Stages of Alzheimer-related neurofibrillary changes were visualized using the Gallyas silver stain [6]. Extracellular deposits of Aβ peptide were immunostained with the mouse anti-β-amyloid 17-24 antibody (1:5000, clone 4G8, BioLegend, Koblenz, Germany). Alpha-synuclein pathology was detected with the anti-syn-1 antibody (1:2000, clone number 42; BD Biosciences, CA, USA). For visualization of microglia and vessels, sections were treated with 10 % methanol and 3 % concentrated H_2_O_2_ in Tris-buffered saline (TBS). Epitopes were unmasked using pretreatment with 1.3 µg/ml proteinase K for 10-15 min at 37 °C (Invitrogen, Darmstadt, Germany). After blocking of unspecific binding sites with bovine serum albumin (BSA), sections were incubated with the primary antibody against IBA1 (1:500, Abcam, Cambridge, UK) over night, a secondary biotinylated antibody (1:200; 2 h, room temperature, Vector Laboratories, Burlingame, CA, USA), and the avidin-biotin-peroxidase complex (ABC Vectastain, Vector Laboratories, Burlingame, CA, USA). The immunoreaction was visualized using 3,3’-diaminobenzidine tetrahydrochloride (DAB; Sigma Taufkirchen, Germany). Next, sections were washed with TBS at 95°C for 5 min, retreated with 10 % methanol and 3 % concentrated H_2_O_2_, and incubated for 48 h with Ulex europaeus lectin I (UEA-l; 1: 800, biotin-coupled, GeneTex, Irvine, CA, USA). Subsequently, sections were incubated with the ABC kit solution, and the reaction product was visualized with a blue chromogen (SK-4700, Vector Laboratories). Omission of all primary antibodies and the lectin resulted in lack of staining.

### Image acquisition and analysis in the SHRSP model

A total of 20 animals were utilized for the immunofluorescence analysis, with 5 SHRSP assigned to early chronic hypertension, 5 SHRSP for late chronic hypertension, and 5 age-matched normotensive controls for each stage of hypertension. Immunofluorescence images were acquired using a Zeiss confocal microscope (LSM 700). The 20x objective was used for overall microglia soma counting, size quantification and blood vessel association. Z-stack images (8-bit, 512 × 512 pixels with a voxel size of 1.25 × 1.25 x 1 µm^3^) were taken from 10 fields of view per animal in the hippocampal CA1 region and overlying retrosplenial cortex, respectively . While a comparatively low hippocampal atrophy has been classically regarded as a parameter that allows the distinction of subcortical cSVD from AD [67], recent evidence suggests that hippocampal atrophy (including subfield CA1) can be associated with cognitive decline in cSVD [9, 33]. In the present study, we therefore focused on hypertension-related microglial changes in the hippocampal CA1 subfields as well as the retrosplenial cortex that is connected to the hippocampal formation through limbic circuits. Single-cell morphological analyses were performed in high-resolution images acquired with the 40x/oil objective. Z-stack images (8-bit, 1024 x 1024 pixels with a voxel size of 0.16 x 0.16 x 0.3 µm^3^) were taken out of 4 fields of view per animal per region, retrosplenial cortex and hippocampal CA1 region, respectively. Thereby, 15-20 microglial cells were analyzed per animal in each brain region. Gain and laser power were kept equal for all animals.

For studying vascular-associated microglial, image analyses were performed using openly available ImageJ software and a Phyton3-based in-house routine. For hippocampal images, rectangles were manually cropped out, so that the CA1 region was included. Pre-processing contained z-projection (maximum intensity), subtraction of background (rolling ball radius: 50 pixels), and median filter to remove salt and pepper noise. For microglia soma segmentation, automatic thresholding with moments dark option and size filtering > 60 µm^2^ was used to quantify the mean microglial soma size of IBA1^+^ cells. For vessel segmentation in STL-labelled images, the OMELETTE framework was adapted [43] (https://gitlab.com/hmattern/omelette). First, blood vessels were enhanced using a multi-scale Frangi filter [17]. To adapt the filter’s vessel sensitivity parameter gamma per image, i.e., to provide robust vessel enhancement for varying pixel intensity distributions, gamma was selected empirically to be 40 % of the image’s maximum absolute Hessian eigenvalues. Thereafter, a multi-scale Frangi filter was applied (filter scales 1, 2, 3, 4, 5 pixels). Subsequently, the blood vessels were segmented from the enhanced images by applying hysteresis thresholding [18]. For hysteresis thresholding, which requires a lower and upper threshold, the three-class Otsu’s method [51] was used to estimate the two required thresholds from the enhancement distribution (self-tuned threshold estimation). Finally, the relative number of microglia soma tangent to the vessel segmentation was estimated. To analyze the three-dimensional (3D) microglia branching morphology in rodents, we applied the 3DMorph automatic analysis workflow via MATLAB on high-resolution images [76]. Using the interactive mode guaranteed correct cell segmentation. Thereby we obtained single cell estimates of cell volume, cell territory volume, i.e., the volume of a 3D convex hull connecting the endpoints of branches, 3D ramification index (i.e. cell volume / territory volume), number of endpoints, number of branchpoints, and average branch length. The analysis was conducted in a blinded manner with respect to the pathological groups.

### Image acquisition in human post-mortem tissue

Images were taken using an Eclipse LV100ND microscope equipped with a digital DS-Fi3 camera and the NIS-Elements software (NIKON GmbH, Düsseldorf, Germany) from the cingulate gyrus of 100 µm-thick hemisphere sections labeled with double-label immunohistochemistry for IBA-1 and UEA-l. The position for image acquisition was randomly selected with the 4x objective and z-stack images were acquired with the 20x objective (image area size 675,470 µm^2^). The N.I.H. ImageJ software was used to generate minimum intensity projections (MIP) from the z-stack images after image brightness was optimized, noise reduction was performed (despeckle command), and the median filter (radius 2.0) was applied in z-stacks. Next, Adobe Photoshop was used for autothresholding and autocontrasting, and to invert the colors of MIP images. The latter step allowed to obtain microglia and vessels with bright colors in a dark background for the quantification of microglia morphologies with established image analyses pipelines. Using ImageJ, the background of inverted images was subtracted (rolling ball radius 50.0 pixels, using „separate colors“ and „sliding paraboloid“ options), and images were reopened in Adobe Photoshop. The inverted blue color of microglia was assigned green and the inverted magenta color of vessels were assigned red using the channel mixer in Adobe Photoshop to further improve the color contrast between microglia and vessels. Inverted images with double labeling were used to quantify the overall microglia cell density and the density of VAMs (analyzed in MIP generated from z-stacks with 25 images; 2880 x 2048 pixels with a voxel size of 0.11 x 0.11 x 1.5 µm^3^). The VAMs were defined as IBA1-positive cells with their somata attached to UEA-l-labeled vessels. To analyze single-cell microglia morphologies, new inverted images were generated (19-53 images in z-stack) to maximize the number of microglia analyzed per case in the MIP images (larger stacks in cases with lower microglia densities, and vic e versa to avoid overcrowding / overlay of cells). From these latter images with IBA1-positive cells (green) and UEA-l-labeled vessels (red), red vessels and shades (out-of-focus areas in MIPs) were removed by using the “remove color” function with thresholding in Adobe Photoshop.

To characterize microglia morphology, the somata were manually outlined for each cell and quantified using the measure function. To investigate the branching complexity of individual cells, microglia were automatically thresholded using the Triangle option and size filtering of >2000 pixels, so that the single-cell branching area as well as the cellular solidity, i.e., cellular branching area / convex hull area, could be estimated. For simplicity and consistency, the cellular solidity is called two-dimensional (2D) ramification index in the rest of this article since it corresponds to the 3D ramification index in rodents. The analyses were performed blinded to the pathological groups.

### Hierarchical Agglomerative Clustering and Uniform Manifold Approximation and Projection (UMAP) for Morphological Features

Single-cell morphology features were standardized by removing the mean and scaling to unit variance. To explore morphological microglia clusters and investigate their latent space in this context, a pipeline for dimensionality reduction was implemented in Phyton 3.9 [44]. Four clusters were extracted using hierarchical agglomerative clustering with Ward-linkage. UMAP analysis was performed for the hippocampal CA1 region and retrosplenial cortex individually using the following UMAP parameters: local connectivity set to 0.5; size of local neighborhood set to 35; minimum distance of two points set to 0.3; Euclidean distance metric used; and 2 components/dimension estimated. To assess the clustering performance, scatter plots in the UMAP-based latent space were generated (color-coded clusters).

### Tissue collection and brain single-cell isolation for FACS analysis

The experimental cohort consisted of a total of 22 rodents, with 5 SHRSP rats assigned to the early hypertension group and their corresponding age-matched normotensive controls, in addition to 6 SHRSP rats assigned to the late hypertension group and their age-matched normotensive controls. Each stage was performed twice. For anesthesia, pentobarbital (40 mg/kg body weight) was intraperitoneally injected in all animals. Transcardial perfusion was conducted with 120 mL of sterile PBS. Brains were divided into two sagittal halves, of which one was processed for flow cytometric analysis, and the other half was processed for microvessels isolation (see further). The hemisphere processed for cell characterization via flow cytometric analysis was cleaned of meninges and dissected into specific regions (cortex and hippocampal region). Each region was collected in a separate tube. Tissues were disrupted in a glass homogenizer in a buffer containing Hanks’ balanced salt solution (HBSS), 1 mM HEPES (pH 7.3), and 45 % glucose and passed through a 40 μm cell strainer. The suspension was centrifuged at 400 g for 20 min, and 1 ml supernatants from each corresponding brain region were transferred into 2 ml polypropylene collection tubes for cytokine immunoassays. The brain suspension was followed by a discontinuous 30 to 70 % (Percoll) density gradient. After myelin aspiration, immune cells were collected from the 30/70 % Percoll interphase, filtered using a 70 μm cell strainer, washed with RPMI, and resuspended in FACS buffer (2 % fetal calf serum in PBS) containing EDTA. Cells were washed, transferred to flat-bottom 96 well plates, and immediately processed for subsequent flow cytometric analysis. According to the cell yield after isolation, each sample was aliquoted three times up to 1.5 x 10^5^ cells adjusted for staining in 100 μl ice-cold FACS buffer. Prior to fluorochrome-conjugated antibody labeling, cells were incubated for 15 min at 4 °C with a purified mouse anti-rat CD32 FcγII (clone D34-485, Rat BD Fc Block) to block unspecific binding. Thereafter, cells were stained with a mixture of fluorochrome-conjugated antibodies in 100 μl of FACS buffer for 30 min at 4 °C. Cells were washed twice and resuspended in 200 μl of FACS buffer. The mixture of fluorochrome-conjugated antibodies included Zombie NIR™, anti-CD45 (clone: OX-1), anti-CD11b (clone: OX-42), anti-CX3CR1 (clone: SA011F11), anti-P2Y12R (polyclonal), anti-CD86 (clone: 24F), anti-CD200R (clone: OX-102), anti-RT1B (clone: OX-6), and anti-CD163 (clone: ED2). Single live cells were identified by the exclusion of doublets, cell debris and gating of viable cells using a live/dead dye (**Suppl. Fig. S3**). Traditionally, microglia are identified as CD11b^+^CD45^int^, whereas CD11b^+^CD45^high^ population corresponds to other central nervous system (CNS) macrophages [21]. However, microglia respond to inflammatory states upregulating CD45 [22], [23], resulting in incorrect classification of microglia as bone marrow-deri ved macrophages. Therefore, in this study, the main populations of cells were identified through Forward Scatter light (FSC) and CD45 expression for both early and late chronic hypertension with their respective age-matched normotensive controls. Target cells were further gated as CD45^+^ and CD11b/c ^+^ cells. Cells positive for CD45, CD11b/c, and P2Y12R were classified as microglia from the hippocampus (**Fig. 2a, b**) and cortex (**Fig. 2e, f**). P2Y12R has been identified as a receptor selectively expressed on microglia and can be used as a marker to distinguish CNS resident microglia from blood-derived myeloid cells [24], [25]. We have developed such gating strategy considering microglia activation appropriate for its senescent and inflammatory response that yields them indistinguishable from recruited myeloid cells. Optimization was performed using antibody titrations, and FMO controls to assess background fluorescence in the respective detection channel. Samples were acquired on AttuneNxT and analyzed with Flowjo Analysis Software (v10.5.3).

### Population identification and high-dimensional data analysis

Sample data were acquired on the AttuneNxT Flow Cytometer (Thermo Fisher Scientific), exported into FlowJo version v10 (TreeStar), compensated, and subjected to unbiased analysis as previously described [8, 39]. First, manually annotated gatings were used to calculate the relative frequencies of microglia and leukocyte populations. Second, microglia manual gating was selected to export live single cells for dimensionality reduction. UMAP analysis was performed for cell visualization from each brain region, i.e., hippocampus and cortex, separately, including exported clean live microglia from both control rats and SHRSP. For the hippocampus: 7,500 microglia cells per animal were randomly subsampled and used to generat e the hippocampus UMAP plot. For the analysis of cortex microglia: 10,000 microglia cells per animal were randomly subsampled and used to generate the combined cortical UMAP plots. A total of 75,000-120,000 cells were analyzed per region and included all compensated markers except dead cells dye staining unless otherwise specified. Microglia cells from both groups were plotted together on a single UMAP for each time point. UMAP parameters were established as follows: local connectivity set to 0.5; size of local neighborhood set to 35; minimum distance of two points set to 0.3; Euclidean distance metric used; and 2 components/dimension estimated. For cluster characterization, the FlowSOM algorithm was run on the merged dataset to cluster every cell after evaluating refined sub-clusters found by Phenograph. Summary tables containing expression levels of each marker, cell frequencies, and cell numbers of UMAP datasets were exported, and total cell fraction per cluster was calculated. Heatmaps display normalized median expression levels from lowest to highest of all markers per merged sub-population group.

### Tissue dissociation for microvascular cell isolation

The brain was divided into two sagittal halves, of which one was processed for FACS (see above), and the other half was processed for microvessels isolation as previously described [52]. The protocol was modified and optimized to obtain pure microvessels derived from the cortices of rat brains. Briefly, cerebral cortices were cleaned of meninges, cerebellum, and brain stem were removed, and the remaining tissue was mechanically reduced into small pieces of ∼1 × 1 mm in size. Homogenate pieces were transferred for digestion into sterile conical tubes with DMEM/F12 containing collagenase type II (1mg/ml), DNase I (15 μg/m), 100 units/ml penicillin, 100 μg/ml streptomycin and 2 mM glutamine. The tissue was mechanically dissociated with a 5 ml pipette and digested in a shaker for 50 min, 37 °C. Myelin and neurons were removed by centrifugation in 20 % (BSA)-DMEM/F12 at 1000 g for 20 min, 4 °C. To maximize brain microvessels retrieval, the resulting myelin layer, together with the BSA supernatant, were carefully removed and underwent two extra centrifugation cycles in the same conditions. Pellets were maintained on ice. After the third centrifugation step, microvessels obtained in the pellet were combined into a new conical sterile tube and further digested with collagenase-dispase (1 mg/ml, Roche) and DNase I (6.7 μg/ml) in DMEM for 60 min, 37°C. In the course of the cell digestion, a 33 % continuous isotonic Percoll density solution was obtained by mixing 1 mL of 10x HBSS, 9 mL of Percoll, 1 mL of 1x HBSS and 1 mL of fetal bovine serum (FBS). The isotonic Percoll mix was then centrifuged (30,000 g, 60 min, 4 °C). The digested homogenate was washed, centrifuged, and overlaid on top of the 33 % continuous isotonic Percoll gradient and centrifuged for 10 min at 1000 g, 4 °C with slow deceleration. Brain microvessels were collected from the corresponding layer and washed twice in ice-cold DMEM. To remove any remaining debris, the cell suspension was transferred through a 40 μm cell strainer for microvessels collection. Thereafter, the cell strainer was reversed and washed into a new tube for the retrieval of microvessels . The cell suspension was centrifuged at 400 g for 10 min at 4 ªC and the purity of the microvessels was quickly evaluated mounting 20 μL of microvessels suspension and observed under an invert ed microscope. The structural features of the microvessels preparation were first examined using a bright field microscope (Primo Star Carl Zeiss, Jena, Germany) and brain debris, myelin and white matter were not observed in the final preparation. Microvessels retained their structure and were isolated to obtain quantitative data representing changes in gene expression and activation of several vascular signaling pathways. The microvessel pellet was resuspended in 200 μl of RNA*later* (AM7020, Thermo Fisher) and stored for further RNA isolation processing.

### RNA isolation from cortical microvessels

Collected microvessels were pelleted at 20,000 g for 10 min and resuspended in 350 μl of RLT Plus Buffer from the RNeasy® Micro Kit (QIAGEN Inc.). Total RNA was isolated using the RNeasy® Micro Kit according to the manufacturer’s instructions. Samples were dissolved in an appropriate amount of RNAase-free ddH2O and RNA concentration and purity were determined using NanoDrop 2000 spectrophotometer (Thermo Fisher) and stored at −80 °C until further use.

### RT-qPCR

Gene expression levels of tight junction proteins and adhesion molecules, were assessed in triplicates using 10 ng isolated RNA. Relative gene expression was determined using the TaqMan® RNA-to-CT™ 1-Step Kit (Thermo Fisher Scientific). Reactions were developed in a LightCycler® 96 (Roche). Reverse transcription was performed for 30 min at 48 °C, followed by inactivation for 10 min at 95 °C. Subsequently, a two-step amplification was run for 55 cycles, comprising of denaturation for 15 s at 95 °C and annealing/elongation for 1 min at 60 °C. TaqMan® Gene Expression Assays used for mRNA amplification are listed in **Suppl. Table S2**. For quantification analysis, the threshold cycle (Ct) using the comparative 2^-ΔΔCt^ method was used [57]. *Gapdh* mRNA expression levels were chosen as reference and relative target gene mRNA levels were determined by the target gene/reference gene ratio. The resulting data were further normalized to values of appropriate control groups and expressed as fold change in arbitrary values.

### Statistical Analysis

All data points were assessed for Gaussian distribution using the Shapiro-Wilk test. For two group comparisons, the statistical significance was analyzed using a two-tailed Student’s *t* test with Welch’s correction for unequal variances. For conducting multiple comparisons in rodents, a two-way analysis of variance (ANOVA) was performed, followed by Holm-Sidák’s post hoc test. Group and age were considered as categorical variables to examine the statistical significance between hypertensive vs. control rats and to identify potential age-related effects. Data analysis was performed using GraphPad Prism software v.9.3.1 (San Diego, CA). The rodent data shown are representative of four independent experiments, two per time point. All quantified values are represented as independent data points and as mean ± SEM unless specified otherwise. *P* values were considered to indicate statistical significance: * for *p* ≤ 0.05; ** for *p* ≤ 0.01; *** for *p* ≤ 0.001; **** for *p* ≤ 0.0001.

### Software

Flowjo UMAP and Phenograph plugins. Flowjo v.10. Installation of R and R libraries: flowCore, FlowSom, pheatmap, matrixStat. Phyton 3.9. Open-source UMAP package via Phyton (https://umap-learn.readthedocs.io/en/latest/index.html). MATLAB-based Cyt3 software. Open-source script 3DMorph via MATLAB. ImageJ: Rasband, W.S., (https://imagej.nih.gov/ij/, 1997-2018).

## Results

### Distinct microglial morphological features in the cortex of patients with cerebral small vesse l disease

We conducted an in-depth investigation into the potential alterations of microglia density and morphological complexity using IBA1^+^ cells from the cingulate cortex of elderly individuals diagnosed with cSVD **(Fig. 1)**. Our analysis revealed a significant increase in microglia density in cSVD tissue samples **(Fig. 1a-e)**. In addition, we quantified the total number of VAMs **(Fig. 1g-i)** and determined their density with respect to the total number of IBA1^+^ cells quantified in a defined area, and found a significant increase of VAMs in cSVD individuals compared to control cases without cSVD **(Fig. 1f)**. Furthermore, morphological analysis of individual microglial cells **(Fig.1c, d)** revealed that IBA1^+^ cells from cSVD cases exhibited a significantly enlarged somatic area compared to controls **(Fig. 1g)** and displayed a decreased 2D ramification index, i.e., microglia branching area / convex hull area, indicating a lower degree of cell ramification and branching complexity **(Fig. 1h, i)**.

**Fig. 1.**
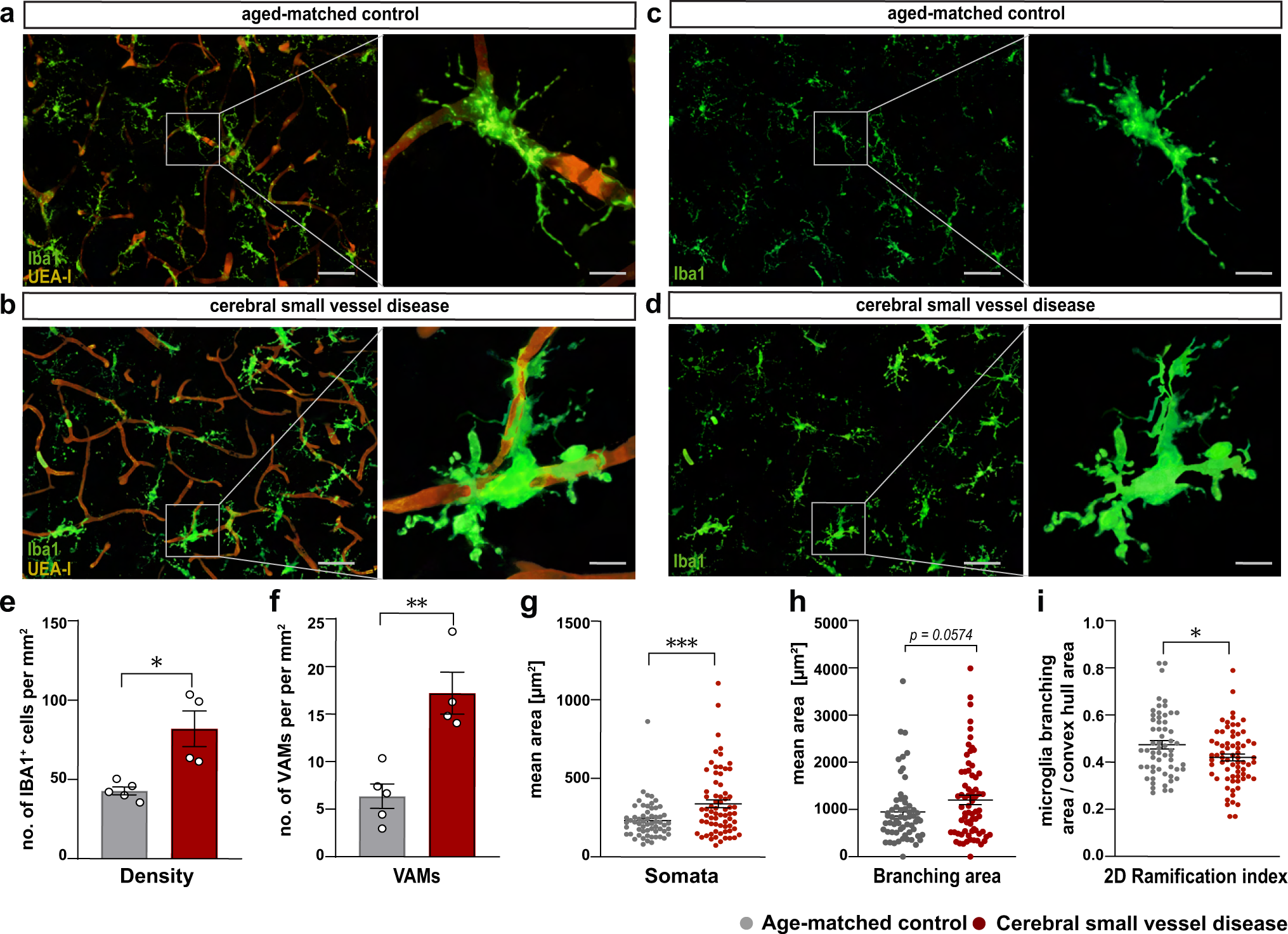
Distinct morphological features of microglia, including enlarged somata thickened processes, and increased territorial volume, observed in post-mortem tissue from patients with cerebral small vessel disease. **(a)** Representative overview and micrographs of individual vessel associated microglia (VAMs) in post-mortem tissue of aged controls and (b) cSVD individuals. (c) Representative micrographs of individual IBA1^+^ microglia of aged controls and (d) cSVD individuals. (e) Quantification of microglia density within a defined area of 1,013,205 µm^3^ standardized to 1mm^2^. (f) Quantification of VAMs within the defined region. (g-i) Quantification of different morphological features of microglia (soma size, average branch area and ramification index). Individual data points indicate averaged microglia data from an individual donor (e, f) or individual microglia within the defined region (g-i). Columns and error bars show mean ± SEM; (a-f), n = 5 controls, n = 4 cSVD subjects. (g-i), n = 145 microglia in age-matched controls, n = 222 microglia in cSVD. *p*-values: * ≤ 0.05; ** for *p* ≤ 0.01; *** for *p* ≤ 0.001; (scale bar overview image = 100 µm; single cell = 25 µm)

### Morphological changes in hippocampal and cortical microglia are already visible in early hypertension in the SHRSP model of cSVD

Animals in the SHRSP group exhibited elevated systolic blood pressure from the age of 8 weeks onwards compared to age-matched controls (8- and 24-week s p < 0.001, 34-weeks p = 0.001, **Suppl. Fig. S1)**. We performed single-cell morphological analyses using high-resolution confocal images to investigate the effect of chronic hypertension on microglial morphology and distribution in the hippocampal CA1 region and retrosplenial cortex **(Fig. 2 and Suppl. Fig. S2)**. Using unsupervised clustering of all available single-cell morphological features, we obtained four different clusters. These clusters reflect a continuous morphological transition, instead of strictly distinct morphological categories **(Fig. 2g-i, p-r)**. Assessing morphological features separately, we observed a significantly increased microglial somatic area, a marker of cell activation, in both brain regions already in early chronic hypertension, which persisted in late chronic stages. Additionally, microglia in late chronic hypertension displayed a significantly more ramified phenotype compared to controls in both brain regions (increased number of endpoints / maximum branch length and decreased 3D ramification index) **(Fig. 2c-f, l-o)**. These results are in line with the morphological alterations found in human analyses and suggest that chronic hypertension induces microglial morphological changes, potentially indicating cellular reactivity, which could contribute to neuroinflammation and neuronal damage in the hippocampus and retrosplenial cortex.

**Fig. 2.**
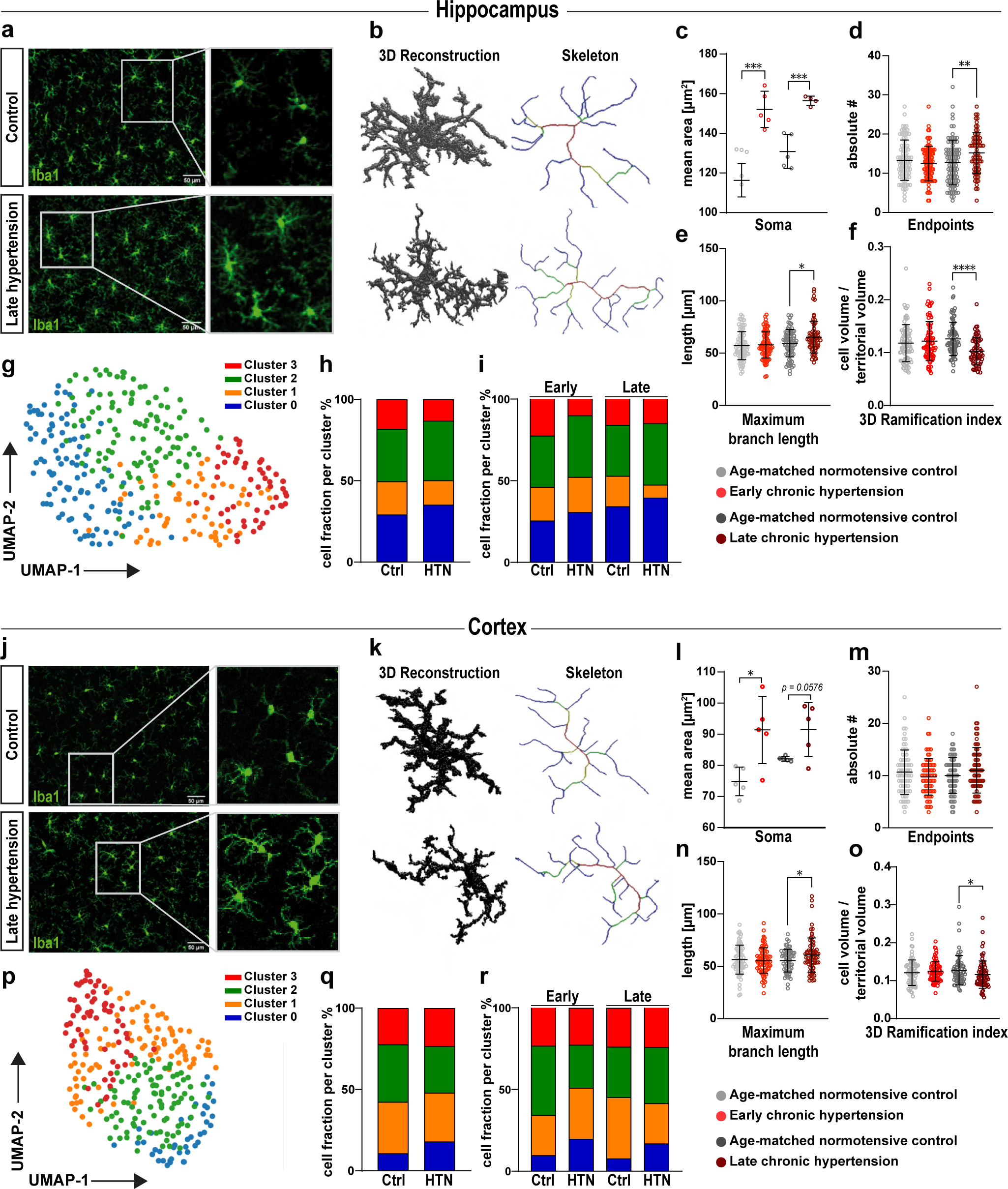
Microglia morphological responses in hippocampal and cortical subfields in chronic hypertensive SHRSP rats. Representative high-resolution images of 15-20 microglia cells of 4 fields of view per animal in the hippocampus **(a)** and in the cortex **(j)**. Confocal three-dimensional reconstructions of IBA1^+^ microglia (green) in late chronic hypertension showing altered skeletal geometrical features and morphological changes induced by chronic hypertension in the hippocampus **(b)** and in the cortex **(k)**. **(c-f)** Quantitative microglial morphological changes derived from 3D microglial reconstructions in the hippocampus and in **(l-o)** the cortex. 15-20 microglial cells were analyzed from 5 independent rodents for each group and brain region. Individual data points indicate averaged microglia data from an individual donor (**c** and **l**; soma area) or individual microglia within a defined region (**d-f** and **m-o**; endpoints, max branch length, and ramification index). **(g)** UMAP plot of microglia clusters based on major morphological features in the hippocampus and **(p)** the cortex. (h) Relative frequencies of microglia per cluster in the hippocampus and **(q)** the cortex. **(i)** Microglia morphological cluster distribution according to the phase of hypertension in the hippocampus and **(r)** the cortex. Data represent mean ± SEM. ANOVA and Holm-Sidák’s post hoc test for multiple comparison were applied to test for significant differences. Ctrl, Controls; HTN, Hypertension. *p*-values: * ≤ 0.05; ** for *p* ≤ 0.01; *** for *p* ≤ 0.001; **** for *p* ≤ 0.0001

### Phenotypical characterization of microglia in chronic hypertensive states

Next, we characterized microglial populations by analyzing their surface marker expression in both early and late chronic hypertensive states by flow cytometry **(Fig. 3a, b and e, f).** Chronic hypertensive states revealed intriguing regional differences at early stage of hypertension, whereas similarities between the hippocampus and cortex were observed in the control groups. Median fluorescence intensity (MFI) for FSC was used to investigate microglial physical properties. In the hippocampus, aging led to a reduction in microglia frequency. Despite the reduction in microglia counts (% of single live cells) observed in the hypertensive hippocampus during late stages of hypertension, a notable enlargement in microglial cell soma size was observed as the duration of hypertension persisted, and this size difference was significant when compared to normotensive hippocampus **(Fig. 3c)**. In the cortex, microglia frequency decreased during aging in both normotensive and hypertensive brains, but chronic hypertension was associated with a higher microglia frequency **(Fig. 3g)**. Moreover, comparable to the findings in the human cSVD cortex **(Fig. 1g)**, chronic hypertension resulted in an increase in microglial cell size in both the hippocampus and cortex. Interestingly, aging led to a decrease in microglia frequency but an increase in cell size and count in the brains of aged control rats when compared to their young counterparts **(Fig. 3c, g)**. Subsequently, we focused our analysis solely on CD45-, CD11b/c-, and P2Y12R-gated microglia to investigate the density of markers associated with anti-inflammatory and pro-inflammatory microglial pathways.

**Fig. 3.**
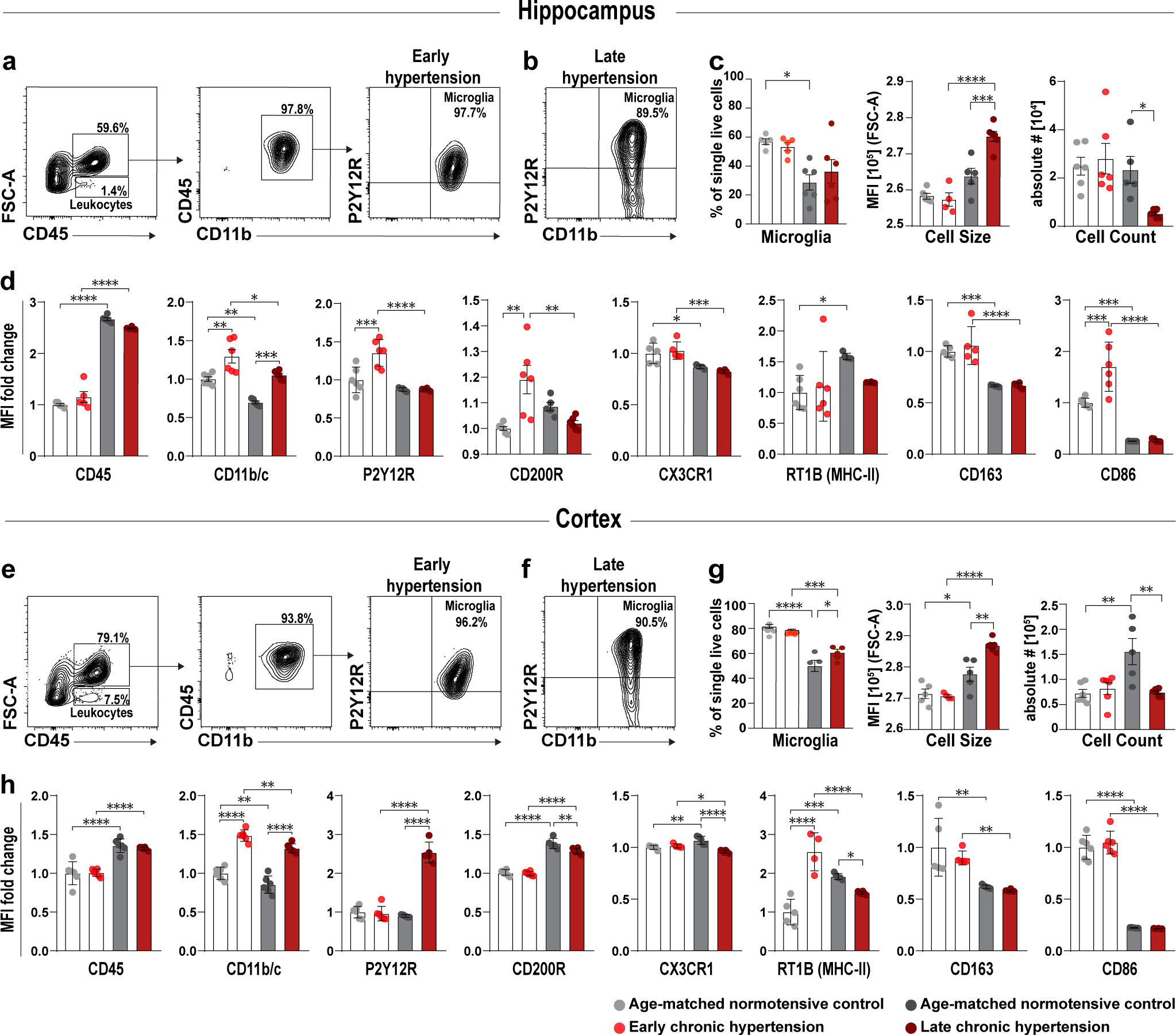
Hippocampal and cortical microglia cell reactivity, age-related effects and dynamic activation in chronic hypertensive states. **(a, e)** Identification of microglia derived from the hippocampus and cortex of SHRSP rats via FACS analysis. Single live cells were identified by the exclusion of doublets, cell debris and gating of viable cells (Suppl. Fig 3). Main populations of cells were identified through Forward Scatter light (FSC) and CD45. Leukocytes were selected based on their size and CD45 expression. Higher FSC CD45^+^ events were further gated as CD45^+^ and CD11b/c ^+^ positive events and further classified by their expression of P2Y12R and CD11b/c. Double positive events were classified as microglia in early chronic hypertension and **(b, f)** late chronic hypertension. **(c)** Frequency and count of microglia cells in the hippocampus and **(g)** the cortex, median fluorescence intensity (MFI) for FSC was used to investigate microglia size. **(d, h)** Bar charts showing the fold change in MFI in each surface antigen investigated in hippocampal and cortical microglia. Individual values shown in the graph represent mean ± SEM. *p*-values: * ≤ 0.05; ** for *p* ≤ 0.01; *** for *p* ≤ 0.001; **** for *p* ≤ 0.0001

### Age-dependent and hypertension-dependent regional differences in microglial surface marker expression profile

To characterize microglia in homeostasis compared to early and chronic hypertension, we compared the expression of distinctive markers that indicate dynamic stages based on the median marker expression of eight proteins (CD45, CD11b/c, P2Y12R, CX3CR1, CD200R, CD163, MHCII (RT1B) and CD86) **(Fig. 3d, h)**. Both hippocampal and cortical microglia undergo an age-dependent increase in CD45 expression, and both hypertensive and normotensive controls exhibit age-dependent decreases in CD11b/c, CX3CR1, CD163, and CD86 surface markers. In early hypertension, hippocampal microglia had increased expression of P2Y12R which was later suppressed, while cortical microglia did not show any significant difference in P2Y12R expression at this stage. CD200R and CD86 were upregulated in early chronic hypertension in hippocampal microglia, whereas cortical microglia did not show any significant difference in these markers. The expression of MHC class II molecules in the CNS was age-dependent in both the hippocampus and the cortex. These differences indicate that hippocampal microglia react earlier and more robustly to chronic hypertension than cortical microglia.

### Phenotypic variations in microglia clusters in the hippocampus

After traditional manual gating analysis of microglia, we applied an additional unbiased multidimensional approach to interrogate the composition and phenotypic variations of microglia subpopulations in the different brain regions. Microglia subpopulations were visualized by UMAP in early **(Fig. 4a-e)** and late chronic hypertension **(Fig. 4f-j)**. This computational approach yields cells of similar phenotypes to be localized into similar coordinates, facilitating the qualitative assessment of different sub-populations inside the microglia population after automated clustering using Phenograph. This approach is helpful for the identification of novel microglia subpopulations that otherwise could be lost if the analysis is performed solely by standard manual gating. Discrete phenotypic variations were identified within the microglia sub-populations, and the co-expression of surface molecules measured on each sub-cluster was plotted into a heatmap for detailed sub-cluster visualization **(Fig. 4d, i)**. Our results revealed the existence of a distinct group of microglial cells expressing specific surface markers (CD86, CD11b/c, and CD163) only in early hypertension, represented by cluster 3 (C3) **(Fig. 4b, c)**. A reduction in the population density of microglial cells within the cluster 2 (C2) was also detected during early stage of hypertension, possibly due to a significant proportion of these cells undergoing transition, as indicated by their allocation to the cluster 3 (C3). Further analysis revealed that the cluster 2 (C2) exhibited an increased expression of P2Y12R and CX3CR1 compared to normotensive microglia **(Fig. 4e)**, suggesting that the upregulation of P2Y12R observed in the overall hippocampal microglial assessment **(Fig. 3d)** was derived from this specific cluster.

**Fig. 4.**
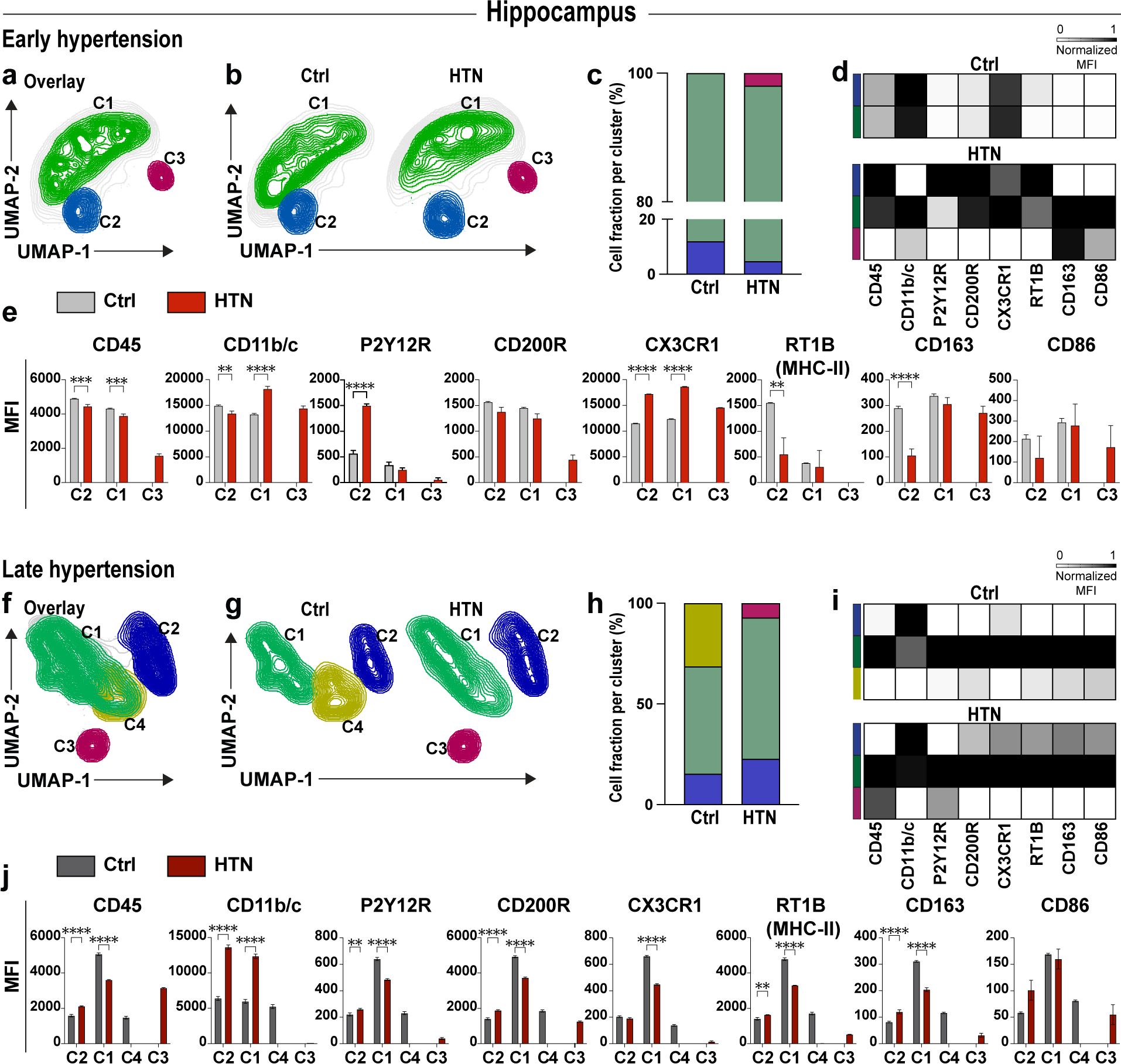
Chronic hypertension results in microglia dynamical changes in the hippocampus of hypertensive rats. Overlay of Uniform manifold approximation and projection (UMAP) map displaying randomly sub-sampled microglia cells from normotensive and hypertensive hippocampus analyzed by flow cytometry in **(a)** early and **(f)** late hypertension. **(b, g)** UMAP plot with color-coded Phenograph-guided clustering with cell identities established based on the investigated surface markers. Representative UMAP plots from normotensive and hypertensive brains displaying microglia sub-clusters corresponding to each cohort. **(c, h)** Relative frequencies of microglial cells and their respective percentages per cluster in early and late chronic hypertension. **(d, i)** Heat map displaying normalized median marker expression values for each population present in each color-coded cluster exhibiting dynamic expression of diverse microglia activation markers (CD200R, CD45, CD86, CD163, RT1B, CD11b/c, P2Y12R, CX3CR1) upon early and late chronic hypertension compared to their respective age-matched normotensive control. **(e, j)** Bar charts showing the median fluorescence intensity (MFI) of each surface antigen investigated in each hippocampal microglia cluster. Bar charts represent mean ± SEM. Ctrl, Controls; HTN, Hypertension. *p*-values: ** for *p* ≤ 0.01; *** for *p* ≤ 0.001; **** for *p* ≤ 0.0001

Aged normotensive controls showed a subpopulation of microglia absent in late hypertension, the cluster 4 (C4), which comprised one-third of the total microglia population. Cells with similar properties cluster together in a 2D UMAP plot, suggesting that this subpopulation may be transitioning from cluster 1 (C1) (homeostatic markers) to cluster 2 (C2) (overall reactivity) **(Fig. 4g-i)**. In late hypertension, a shift in the microglia population from the predominant cluster 1 (C1) to a higher fraction in the cluster 2 (C2) was detected. This shift was accompanied by a decrease in the expression of CD200R, CD45, CD163, RT1B (MHC-II), P2Y12R, and CX3CR1 in the cluster 1 (C1) and an increase in CD11b/c expression. Additionally, cluster 2 (C2) showed significantly higher expression of CD200R, CD45, CD163, RT1B (MHC-II), CD11b/c, and P2Y12R, indicating a second reactive microglial cluster. These results suggest that microglia from cluster 1 (C1) were transitioning to cluster 2 (C2) with increased reactivity in late hypertension. Furthermore, the cluster 3 (C3) expressed a unique combination of CD45, CD86, and P2Y12R, which was not observed in normotensive controls. Overall, our automated clustering approach revealed discrete phenotypic variations and different co-expression patterns of surface molecules at early and late stages of hypertension. In the hippocampus, the transition from early to late hypertension was characterized by distinct phenotypic variations in microglia cluster 1 (C1), including a shift in the density of microglia predominantly located in cluster 1 (C1) to an increased fraction in the cluster 2 (C2) in late hypertension, which indicated the presence of a second reactive microglial subpopulation with heightened reactivity and altered marker expression.

### Phenotypic variations in microglia sub-populations in the cortex

Unsupervised graph-bas ed clustering (PhenoGraph) of cortical microglia cells in early **(Fig. 5a-e)** and in late chronic hypertension **(Fig. 5f-j)** created a 2D map of microglia sub-populations within groups. Analysis of microglia during early hypertension revealed six clusters in normotensive brains, while five clusters were observed in hypertensive brains. During late chronic hypertension, five clusters were observed in normotensive brains, while seven clusters were observed in hypertensive brains. These distinct sub-populations of microglia were identified based on their specific patterns of expression of pro and anti-inflammatory pathways. Further examination of these sub-populations revealed that the highest fraction of microglia localized similarly between groups in cluster 1 (C1). Notably, some clusters were absent or unique in one group only. For instance, microglia in cluster 5 (C5) (**Fig. 5a-c)** that expressed CD45, CD11b/c, P2Y12R, CD200R, and RT1B (MHC-II) (**Fig. 5d, e)** were entirely absent in hypertensive brains, whereas cluster 2 (C2) was present in both groups, with a greater proportion of microglia localized within this cluster in hypertensive brains. Moreover, the cluster 3 (C3) observed in hypertensive brains displayed heightened expression of CD200R and CX3CR1 but lower expression of CD45 and CD86 **(Fig. 5e)**. On the other hand, the cluster 4 (C4) seen in hypertensive brains expressed higher levels of CD45 and CD11b/c **(Fig. 5e)**.

**Fig. 5.**
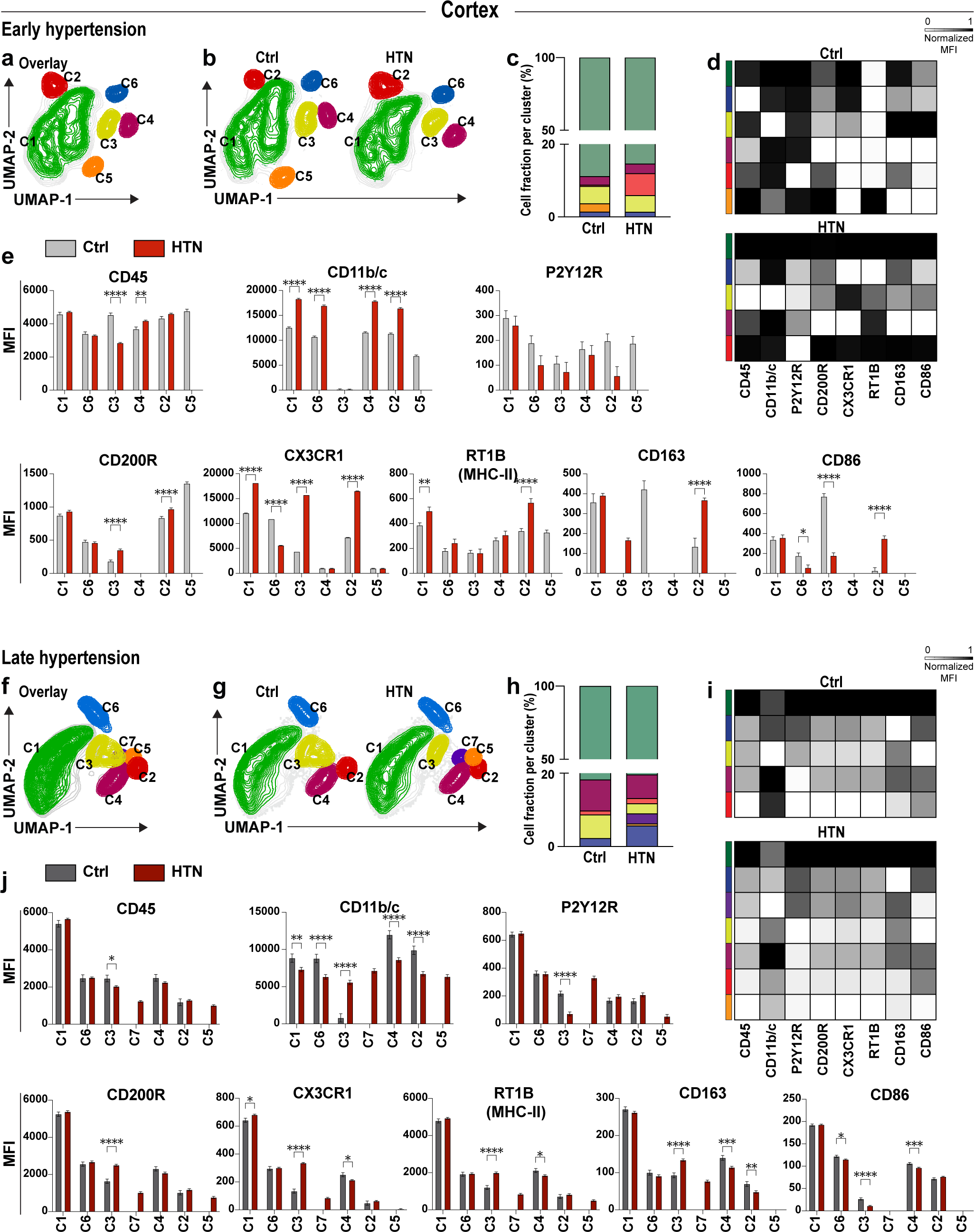
Chronic hypertension results in microglia dynamical changes in the cortex of hypertensive rats. Overlay of Uniform manifold approximation and projection (UMAP) map displaying randomly sub-sampled microglia cells from normotensive and hypertensive cortex microglia analyzed by flow cytometry in **(a)** early and **(f)** late hypertension. **(b, g)** Representative UMAP plots from normotensive and hypertensive brains displaying microglia sub-clusters corresponding to each cohort. (c, h) Relative frequencies of microglia cells and their respective percentages per cluster in early and late chronic hypertension. (d, i) Heat map displaying normalized median marker expression values for each population present in each color-coded cluster exhibiting dynamic expression of diverse microglia activation markers (CD200R, CD45, CD86, CD163, RT1B, CD11b/c, P2Y12R, CX3CR1) upon early and late chronic hypertension compared to their respective age-matched normotensive control. (e, j) Bar charts showing the median fluorescence intensity (MFI) of each surface antigen investigated in each hippocampal microglia cluster. Bar charts represent mean ± SEM. Ctrl, Controls; HTN, Hypertension. *p*-values: * ≤ 0.05; ** for *p* ≤ 0.01; *** for *p* ≤ 0.001; **** for *p* ≤ 0.0001

Subsequent analysis indicated that during late chronic hypertension **(Fig. 5f-h)**, microglia were again primarily localized in cluster 1 (C1). While a lower fraction of hypertensive microglia was found in cluster 3 (C3), these cells displayed higher reactivity with increased expression of CD200R, CX3CR1, CD163, RT1B (MHC-II) and CD11b/c **(Fig. 5i, j)**. Whereas CD200R and CX3CR1 are anti-inflammatory markers, CD163, RT1B (MHC-II) and CD11b/c are associated with phagocytic and antigen-presenting functions that might indicate a state of activation and/or an attempt to suppress pro-inflammatory pathways. Additionally, two clusters of microglia (C5 and C7) were identified, which were absent in normotensive controls. The cluster 7 (C7) expressed high levels of CD200R, CD45, CD163, RT1B (MHC-II), CD11b/c, P2Y12R, and CX3CR1, while the cluster 5 (C5) expressed CD200R, CD45 and CD11b/c at the same level as cluster 7 (C7), but with lower expression of RT1B (MHC-II) and P2Y12R. The phenotypic variations in microglia clusters observed in the cortex during early and late chronic hypertension exhibited greater complexity compared to the hippocampus, as evidenced by the presence of a larger number of distinct clusters. While both regions displayed unique cluster compositions, the cortex demonstrated a more intricate constellation of sub-populations, with specific clusters absent or unique to hypertensive brains. However, the dynamic changes observed in surface marker patterns during the progression from early to late hypertension demonstrate that the predominant density of microglia in the cluster 1 (C1) remained relatively consistent, implying that certain sub-populations arise specifically due to hypertension-induced changes while others are associated to the aging process.

### Microvascular pathology and immune cell infiltration in hypertension-induced BBB disruption

To investigate temporal changes in BBB integrity and the correlation of BBB disturbances with microglial responses in chronic hypertension, we examined isolated fragments of microvessels and found a significant downregulation of the two critical tight junction molecules claudin-5 (*Cldn5*) and occludin (*Ocln*) in early hypertension, indicating impaired BBB integrity as an early chronic hypertension signature **(Fig. 6a-c)**. However, no disturbance in BBB permissiveness was observed in the late chronic hypertensive phase, suggesting compensatory or adaptive mechanisms to repair the BBB and maintain its function in the face of chronic hypertension-induced microvascular pathology. Vascular cell adhesion molecule-1 (*Vcam1*) and intercellular adhesion molecule-1 (*Icam1*) were upregulated with increasing age **(Fig. 6d, e).** An increase in the frequency of leukocytes within the cortex and hippocampus during the early phase of hypertension was detected, indicating potential BBB leaks of these cells from the blood stream, which is in line with the downregulated tight junction molecules, an impaired BBB and potentially neuroinflammatory responses. These findings suggest that hypertension-induced disturbances in BBB integrity may contribute to an increase in leukocyte infiltration, which may vary depending on the stage of the disease and may be further exacerbated by the aging process **(Fig. 6f, g)**. Additionally, higher frequencies of VAMs were present in both the hippocampus and cortex in early and late chronic hypertension, similarly as in the human analysis, suggesting the potential involvement of VAMs in vascular (dys)function and associated pathologies pointing towards a role of microglia in BBB dynamics. Our results suggest a biphasic course of hypertension-induced BBB disruption, immune cell infiltration, and VAM reactivity in the brain and highlight the complex interplay between microvascular pathology and microglial responses in the context of chronic hypertension **(Fig. 6h-k)**.

**Fig. 6.**
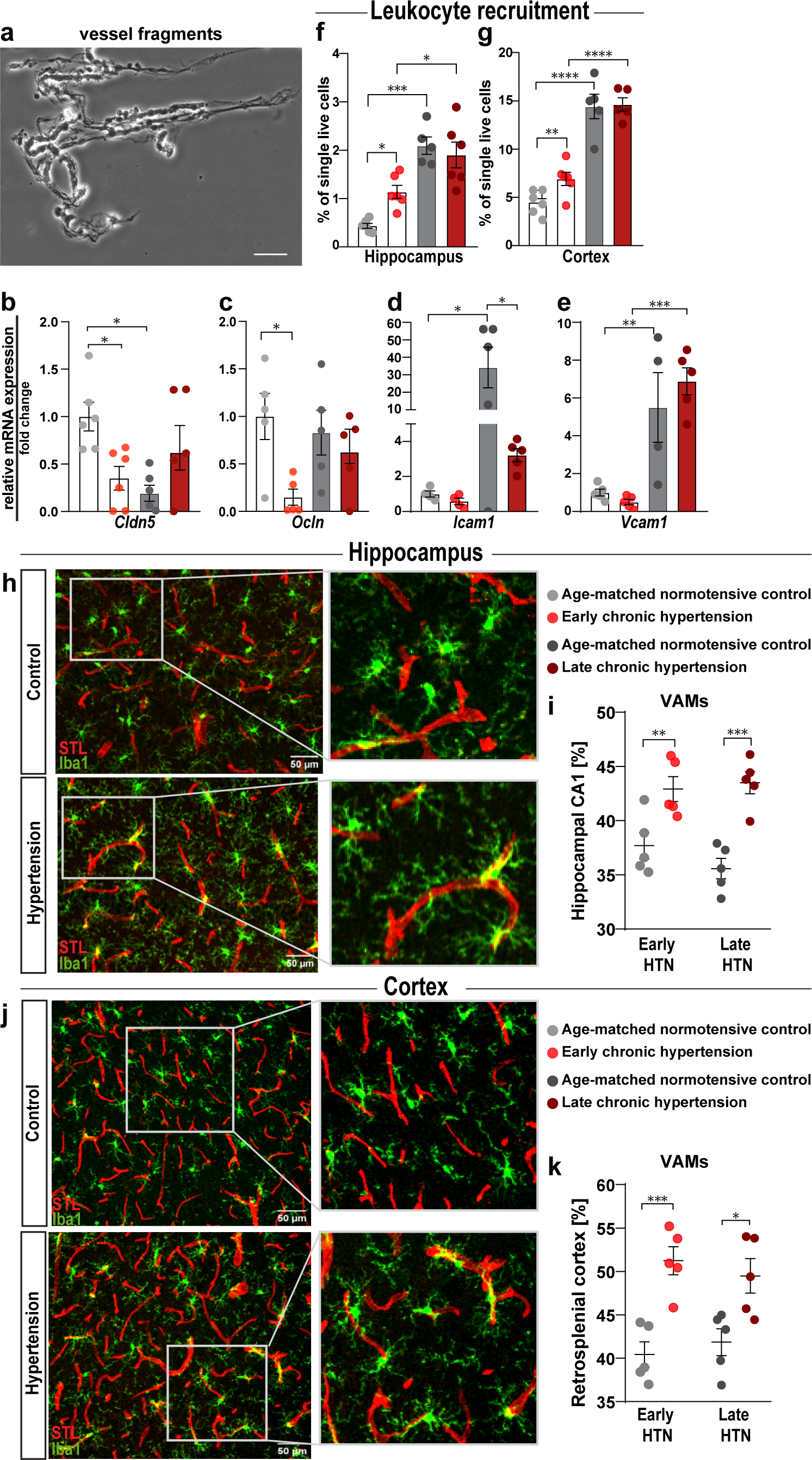
Converging mechanisms of cerebrovascular remodeling and microglia activation in chronic hypertensive states with BBB disruption and leukocyte infiltration into the CNS, and the emergence of vascular-associated microglia. **(a)** Representative phase-contrast image of isolated microvessel derived from brain cortices plated on a microscope slide (scale bar, 50 µm). Quantification changes in gene expression in cerebral microvessels of tight junction genes (b) *Cldn5*, (c) *Ocln,* and adhesion molecules (d) *Icam1* and (e) *Vcam1* at early and late chronic hypertension shown by qRT-PCR. in normotensive and hypertensive rat brains. Fold changes were calculated by normalizing gene expression levels to GAPDH. The resulting data were further normalized on mean values of normotensive controls. **(f)** Recruitment of leukocytes into the hippocampus and cortex **(g)** of all groups in early and late chronic hypertension, represented as the frequency of low FSC CD45^+^ events from the total single live cells. **(h)** Immunofluorescence confocal microscope image of microglia stained with anti-IBA1 representative from hippocampal CA1 region of late hypertensive rats (lower) and its age-matched normotensive control (upper). **(i)** Frequency of vascular-associated microglia (VAMs) in the hippocampal CA1 region. **(j)** Immunofluorescence microglia staining representative from retrosplenial cortex with anti-IBA1 of late hypertensive rats (lower) and its age-matched normotensive control (upper) **(k)** Frequency of VAMs in the retrosplenial cortex. BBB, blood-brain barrier; CNS, central nervous system; HTN, Hypertension; IBA1, ionized calcium-binding adapter molecule 1 (microglial marker); STL, solanum tuberosum lectin-fluorescein isothiocyanate (endothelial marker). *p*-values: * ≤ 0.05; ** for *p* ≤ 0.01; *** for *p* ≤ 0.001; **** for *p* ≤ 0.0001

### Correlations between morphological alterations in microglia and their phenotypic marker expression profiles in the aging brain and in chronic hypertensive states

We assessed the correlation between the expression profile of microglial surface markers and morphological features of microglia in the cortex and hippocampus of normotensive and hypertensive brains **(Fig. 7).** In the hypertensive hippocampus, early chronic hypertension was associated with positive correlations between CD11b/c and RT1B (MHC-II) expression, suggesting similar microglial activation and antigen presentation. Additionally, a positive correlation between CD200R and P2Y12R expression may indicate a potential neuroprotective role for microglia. In contrast, the negative correlation between CD11b/c and P2Y12R suggested a phenotypic shift from surveillance to a reactive state in the hypertensive hippocampus **(Fig. 7b)**. During late chronic hypertension, the positive correlation between microglial soma size and P2Y12R expression, may reflect microglial hypertrophy and increased chemotaxis, while the negative correlation between CD11b/c and CD200R is suggestive of an interplay between these two markers in regulating microglial activation in the hippocampus **(Fig. 7d)**. In the normotensive aging hippocampus, positive correlations between CD45 and CX3CR1, and between RT1B (MHC-II) and microglial endpoints, may reflect compensatory mechanisms of antigen presentation and dynamics of microglial process, allowing microglia to actively sense and respond to changes in their environment. Moreover, we found a negative correlation between RT1B (MHC-II) expression and microglial branch length, which may further suggest a tradeoff between antigen presentation and process elongation **(Fig. 7c)**. During the early stage of hypertension, we observed negative correlations between CD45 and CD200R, and between CD11b/c and CD163 in the cortex, suggesting the involvement of CD45 in downregulation of CD200R signaling pathways and classical activation of microglia. Like in the hippocampus, we found a positive correlation between P2Y12R and microglial endpoints, and between P2Y12R and branch points in the cortex that may further indicate the role of this marker in microglial surveillance, phagocytosis, and morphological plasticity **(Fig. 7f)**. At the late stage of hypertension, we observed a negative correlation between microglial soma size and CD163 in the cortex indicative for a shift towards morphological features that facilitates microglial surveillance contributing to tissue repair. The positive correlations observed in the cortex between CD45 and CX3CR1, and between CD200R and RT1B (MHC-II), CD200R and CX3CR1, and CD200R and CD86, implicated a role in microglial reactivity and interactions with neurons **(Fig. 7h)**. During normotensive aging, we found negative correlations in the cortex between CD11b/c and microglial soma size, and between RT1B (MHC-II) and P2Y12R, which suggested smaller microglia soma area and reduced involvement in antigen presentation and surveillance. In addition, we observed positive correlations between microglial soma size and RT1B (MHC-II) and between microglial soma size and P2Y12R that indicated specialization in antigen presentation and phagocytosis, and the positive correlation between P2Y12R and CD86 implied a role surveillance promoting T-cell activation. Positive correlations between CD163 and CX3CR1, and between CX3CR1 and CD86 suggested microglial involvement in anti-inflammatory responses and tissue repair in the aging cortex **(Fig. 7g)**.

**Fig. 7.**
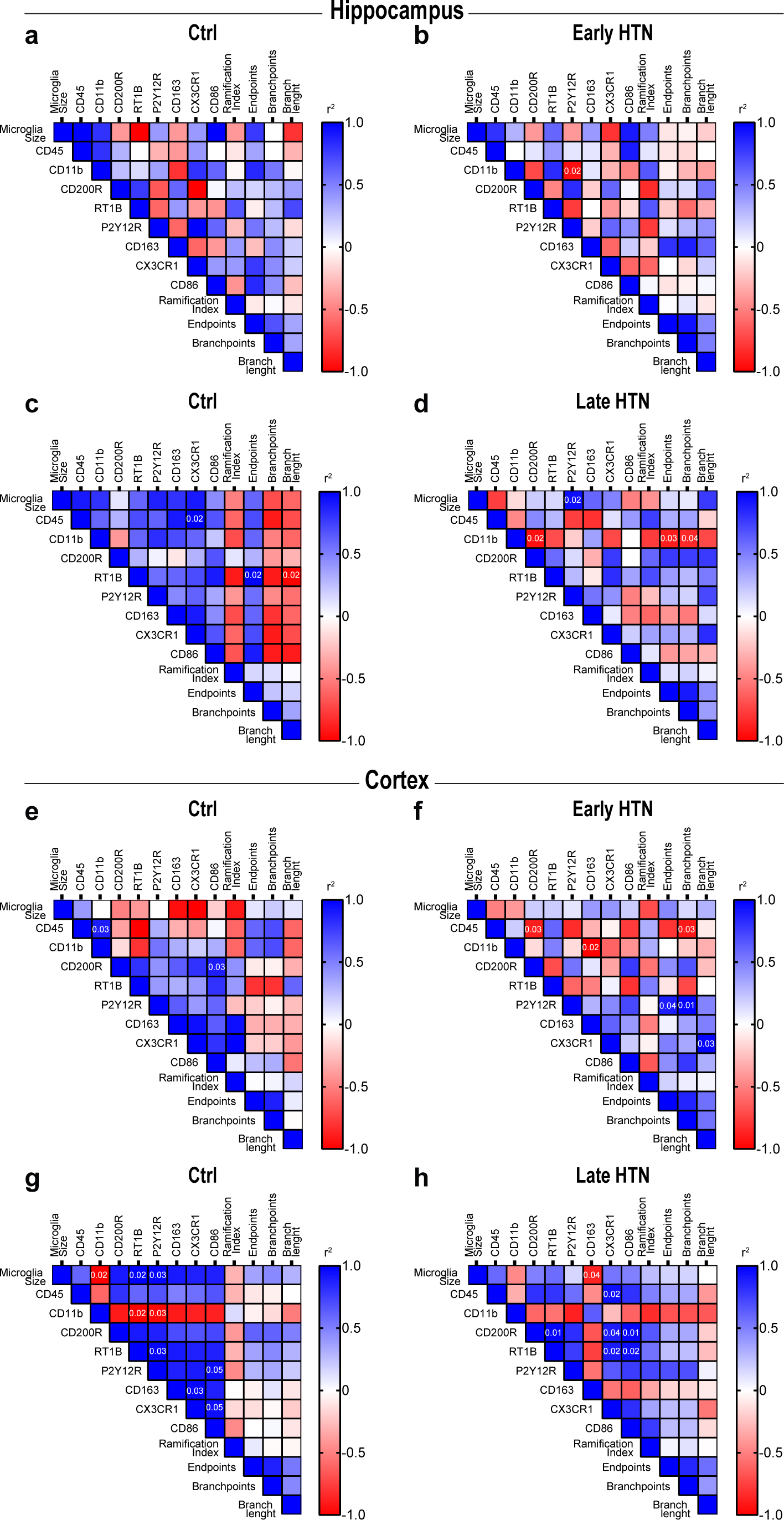
Hippocampal and cortical microglia size and activation markers correlate differently during hypertensive stages. Correlation plots from hippocampal and cortical microglia in **(a, e)** young controls, **(b, f)** early hypertension, **(c, g)** older controls and **(d, h)** late hypertension. Statistically significant values (non-parametric Spearman’s test) are shown in their respective cells. The color of each cell represent s an estimation of the positive (red) or negative (blue) correlation index based on the r value. Ctrl, Controls; HTN, Hypertension

## Discussion

Our study presents distinct features of microglia in human post-mortem cortical tissue of patients with pathologically confirmed cSVD. Specifically, we detected a higher abundance of microglia with thicker processes and with enlarged somata, an increased ramification complexity and a higher association with the vasculature. Expanding our investigation to a rodent model of hypertensive cSVD, we confirmed that the enlarged somata and microglial association to the vasculature were morphological features present already at the early stages of hypertension, whereas the increased ramification complexity appeared during the late stage of hypertension. In a next step, we performed deep phenotypical characterization of microglia surface markers to identify functional subpopulations within the hippocampus and cortex. The dynamic changes detected highlighted early microglial dysregulation in the hippocampus whereas most molecular changes in the cortex occurred firstly at late chronic stages, possibly indicating differences in the regional vulnerability to vascular perturbations. Our findings have an important implication for vascular and neurodegenerative diseases, as the higher vulnerability of the hippocampus suggests a potential involvement in the progression of cognitive decline and dementia associated with cSVD. By identifying distinct microglial morphological changes our results provide a promising basis for further biomarker-based research.

### Microglial morphological changes in cSVD

We presented close similarities between human and rodent microglial morphological changes. Our results suggest that changes in microglial morphology, characterized by enlarged somata and increased ramification, indicate a shift in microglial function [23, 30, 45]. The observed enlargement of microglial soma size and elongated, thickened processes in cSVD cases suggest a reactive response of microglia and potential alterations in their metabolic activity. In a recent autopsy study that focused on the white matter, the authors reported contrasting results to our morphological findings, revealing ameboid microglia in response to hypertension [59]. This observation emphasizes the potential regional difference between white and grey matter on microglial reactivity. To our understanding, no study has investigated microglia morphology in human grey matter during hypertensive cSVD. Different rodent models of hypertension consistently showed enlargement in microglial somata, accompanied by increased microglia ramification in grey matter tissue [29]. These morphological changes were associated with higher levels of CD11b, a marker of microglia reactivity, and even showed the potential reversal of these microglial changes by antihypertensive drugs such as amlodipine [32]. Morphological changes may reflect the impaired ability of microglia to efficiently carry out phagocytosis or to handle the increased load of debris and cellular waste resulting from chronic hypertension and vascular damage. Additionally, their results showed an increase in the phagolysosomal marker CD68 in microglia from hypertensive mice. It is important to note that an elevation of the CD68 marker does not necessarily imply a direct enhancement of phagocytic functions. Instead, it could indicate either an increased engulfment of lysosomal structures or a decreased clearance rate, thereby slowing down microglial catabolism. Consequently, it is conceivable that the observed increase in soma size and thickened processes during chronic hypertensive states might be attributed to potential metabolic alterations, resulting in the distinctive morphological features observed.

### Revealing the vulnerability of the hippocampus in hypertensive cSVD

Evidence suggests that cognitive decline in cSVD is characterized by the coexistence of memory disturbances and cortical symptoms, such as executive dysfunction and attention deficits [20]. A recent investigation revealed an age-related disruption of the BBB in the hippocampus as an early event in the aging human brain [48]. This early BBB leakage observed in mild cognitive impairment, along the AD spectrum, may serve as a relevant starting point to understand similar pathophysiological processes in hypertensive cSVD. Additionally, cardiovascular risk factors impact memory function through WMH [38], potentially enhance the vulnerability of different brain regions to the combined effect of age and hypertension [60] and specifically speeds up neurodegeneration of brain regions, which are connected to WMH. Therefore, it is plausible to hypothesize that hypertensive cSVD exhibits a similar initial occurrence. Comprehensively exploring the potential convergence between hypertensive cSVD and hippocampal vulnerability provide valuable understanding into the specific impact of hypertension on memory function and cognitive impairment. The differences in cell size between control and early hypertension are consistent with the 3D ramification index and the number of endpoints observed in the morphological analysis. Notably, the degree of significance and change observed aligns with the 3D morphological results, which appears to be more robust in the hippocampus compared to the cortex. We hypothesized that chronic microglia adaptation takes place throughout the initiation and progression of the disease, unlike the sudden brain injury that occurs in acute brain injury. Surprisingly, we found that even in early hypertensive states resembling mid-life human chronic hypertension, hippocampal microglia portrayed a highly dynamic heterogeneous phenotype. Notably, at the early hypertens ive stage microglia displayed a pro-inflammatory phenotype evidenced by upregulation of CD11b/c and CD86 and potentially an enhanced chemotactic and anti-inflammatory responsivity as reflected by P2Y12R and CD200R upregulation. Elevated expression of P2Y12R potentially indicated migration of microglia towards injury sites and their preferred localization at the capillary wall. In contrast, upregulation of CD200R suggests activation of anti-inflammatory pathways, potentially serving to mitigate inflammation [63] and potentially counteracting early BBB leaks. The notable increase in CD86, a co-stimulatory membrane receptor, implies microglial priming or transition towards an activated inflammatory state even at early hypertension in the hippocampus. Conversely, the decrease in CD163 expression in both the normotensive and chronically hypertensive aged rats suggests a decline in the anti-inflammatory capability of microglia, confirming that age and hypertension separately affect the marker expression profile of hippocampal microglia, potentially contributing to microglial reactivity and inflammation.

Using a computational approach to identify discrete subpopulations of microglia, we revealed the existence of a distinct group of microglia cells (C3) expressing high levels of specific proteins (CD163 and CD86) in early hypertension that were not present in normotensive controls. Late-stage hypertension seems to result in inefficient regulation of pro-inflammatory conditions, reflected by the suppression of CD200R expression. This suppression is particularly concerning as CD200R signaling modulates immune responses, controls myeloid cell function, inhibits pro-inflammatory cytokine expression, and prevents tissue damage by myeloid-derived cells [40]. We also identified a subpopulation of microglia that were absent in late hypertension but present in aged normotensive controls, and therefore may be transitioning from homeostatic to reactive. Moreover, a subpopulation of microglia showed a higher expression of most markers at the late hypertensive stage, indicating a second reactive and heterogenic microglia sub-population, which differs from the reactive population found upon normotensive aging. This subpopulation may be involved in the activation of pro-inflammatory signaling pathways that lead to the release of cytokines, chemokines, and reactive oxygen species, and therefore may further exacerbate the hypertension-induced neuroinflammatory response.

### Age-dependent and pathological changes in cortical microglia marker expression profile suggest microglia reactivity and primed states in hypertensive rats

Similar to hippocampal microglia, our study detected age-dependent changes in marker expression profile in cortical microglia of hypertensive rats. We observed that the inhibitory immune receptor CD200R was upregulated with aging but significantly lower in hypertensive brains. Interestingly, RT1B (MHC-II) expression was higher in early hypertension, which may precede its final differentiation into disease-associated microglia, identified in different diseases through single-cell RNA-seq [42] but now in chronic hypertension, which is relevant to understand the differential adaptations in microglia, potentially paving the way for disease susceptibility later in life. Our findings demonstrate that during the late phase of chronic hypertension, microglia in the hypertensive cortex displayed a reduced expression of the inhibitory receptor CD200R in comparison to normotensive brains. This reduction may result from hypometabolism due to long-term and constant microglia activation and chronic neuroinflammation and it may also contribute to an inefficient regulation of pro-inflammatory conditions commonly observed in the aging brain, ultimately leading to deficient feedback loops in the induction of neuroinflammatory mechanisms that contribute to neurodegeneration [11, 47, 78]. Moreover, these results suggest there is dysfunction in the CD200-CD200R system, which is necessary for the activation of anti-inflammatory signals supported by previous findings where a significant decrease in CD200R mRNA expression by microglia was found in brains of AD patients [69]. Although most markers display a complex co-expression across clusters, our goal was to characterize the dynamic changes in microglia sub-populations during chronic hypertensive states, in addition to its known progressive senescent modifications, and to provide further evidence of the diversity within the microglia population that is dependent on the stage of hypertension and the CNS region. While both the hippocampus and cortex exhibit distinctive microglia cluster compositions, the larger size of the cortex suggests a greater spatial and functional complexity, potentially requiring a more diverse array of microglia subpopulations. Spatial proteomics analysis or multiple-target immunohistochemistry can be a promising approach for future research, enabling a deeper understanding of the region-specific microglial adaptations and their implications for neuroinflammatory responses during chronic hypertensive states.

### Microglia heterogeneity in chronic hypertension and its implication for BBB leakage and cerebrovascular remodeling

Microglia, known for their heterogeneity, exhibit significant diversity in early development, which decreases in adulthood, resulting in a more homogenous population in steady states [21, 55]. However, during pathological conditions, heterogeneity seems to reemerge, as found here in the cortex and hippocampus in chronic hypertensive states. We observed that microglia in chronic hypertension display a range of morphological and functional characteristics, with overexpression of P2Y12R, a purinergic receptor that regulates microglia activation and migration, indicating a shift towards a pro-inflammatory microglial phenotype and increased reactivity. Our results suggest an overall increase in the chemotactic ability of microglia towards injury, and the clustering of P2Y12R in microglial filopodia may impact their phagocytic activity. Microglia can exist in various reactive states, including their primed state and senescence. Our analysis identified two distinct microglia cell states associated with chronic hypertension, with a significant fraction of surface molecules upregulated in late-phase hypertension that are already increased in early hypertensive states. Microglia that exhibit pro-inflammatory characteristics are known to undergo morphological changes and secrete cytokines, which can cause significant damage to both the vasculature and neuronal tissue, resulting from prolonged exposure to pro-inflammatory mediators [62, 65]. Our findings demonstrate alterations in microglial filopodia, which may impact their phagocytic activity and that can be a result of prolonged exposure to pro-inflammatory mediators, which contribute to the inefficient regulation of inflammation observed in late-phase hypertension. The clustering of P2Y12R identified in our analysis suggests that microglial chemotaxis towards injury may be impacted, potentially contributing to cerebrovascular remodeling and BBB leakage seen in chronic hypertension. These results highlight the presence of a specific subtype of pro-inflammatory microglia during chronic hypertension that can harm brain function and contribute to disease onset by causing persistent tissue damage. The resulting reactive microglia phenotype observed in early hypertension may be a cause or a consequence of the significant disturbances seen in the BBB. In one way, microglia expression levels of purinergic receptor P2Y12R observed in chronic hypertensive states may trigger its migration to vascular sites and lead to a “classical” microglia reactivity pattern, modulating its cytoskeletal network for phagocytosis, followed by remodeling of the vessel wall and significant BBB disturbance, with consequent ingress of leukocytes into the CNS perpetuating microglia activation [46]. Conversely, increased leukocyte adhesion and remodeling of the vessel wall due to chronic hypertension may initiate a microglial “resolving” activation pattern aiding in the resolution of the BBB disturbance that leads to a phase of BBB healing.

### Implications of altered microglial morphological features in cSVD pathogenesis

The decreased ramification index in cSVD microglia, as well as the positive correlation between P2Y12R and microglial endpoints found in the cortex of hypertensive rodents during early hypertension, may indicate a compensatory mechanism in response to chronic hypertension and small vessel damage. P2Y12R is a marker of microglial activation and is involved in process extension and surveying of the microenvironment in an attempt to maintain homeostasis. The overall density of IBA1-positive cells was increased in human cSVD cases compared to controls, although microglia densities were similar to controls in the rodent cSVD model, but our investigation identified a notable presence of microglia with thickened processes in the rodent cSVD group. Notably, we further observed a significant increase in the number of IBA1^+^ cells in contact with blood vessels in both human and rodent cSVD cases, which suggests an augmented density of VAMs. Additionally, hypertension was found to be associated with a significant increase in the fraction of VAMs, even in mid-life hypertensive rats, suggesting early regional neurovascular alterations surrounding blood vessel walls in the disease. Reactive microglia have the potential to exacerbate vascular injury by engaging in excessive pruning, resulting in perivascular cell reactivity and removal of astrocytic end-feet. This can lead to a reduction in the density of Aquaporin 4, which plays a critical role in glymphatic clearance and the removal of protein deposits surrounding the neurovascular unit. In the context of late-stage hypertension, the density of VAMs remained elevated compared to normotensive controls. This suggests that microglia subpopulations may play a dynamic spatio-temporal role over time. During early stages of hypertension, they polarize towards a pro-inflammatory phenotype. As hypertension progresses, there is a persistent increase in anti-inflammatory clusters that could reflect a compensatory mechanism to reduce local neuroinflammation, which if not reduced, it can lead to chronic microglia activation ultimately resulting in self-perpetuating damage to both vascular and neural tissue. Furthermore, microglia in cSVD patients displayed thickened processes, and a significant increase in the soma and the territory they covered, suggesting more complex microglia structures. These findings suggest that microglia in cSVD patients may have undergone changes in their activation state or response to stimulus, potentially indicating their involvement in the pathogenesis of cSVD.

It is now accepted that neuroinflammation plays a significant role in the development of neurodegenerative diseases [24, 37] and alongside neuroinflammation, immunosenescence declines the immune system functions that gradually contribute to critical outcomes such as dementia. Evidence shows that preexisting hypertension exacerbates the development of secondary neurodegenerat ion beyond its acute effect on neurovascular injury [15, 68]. These findings highlight the importance of how managing midlife hypertension is crucial for preserving vascular and brain health. Previously reported findings have indicated that brain tissue surrounding blood vessels in patients with cSVD is affected even before the emergence of lesions such as microbleeds or lacunar strokes [1, 28, 70]. These changes in the extracellular matrix around small brain capillaries may lead to a loss of vascular contractibility, resulting in an inability to meet the high metabolic demands of the aging brain. Regulating vascular risk factors earlier in life may prevent cognitive decline and maintain brain health homeostasis in older age. Our results shed light on the role of microglia in chronic hypertension, suggesting that a subpopulation of microglia may aid in vascular wall maintenance and play a critical role in the neurovascular unit during chronic hypertension. We demonstrated that the presence of chronic hypertension induced morphological changes in microglia, altered BBB permissiveness at an early stage, and promoted neuroinflammation, potentially priming the brain for injury. Our study also revealed that, with late-stage chronic hypertension, a higher percentage of microglia were present in close proximity to blood vessels, suggesting a dynamic spectral role of microglia to enhance neuroinflammatory signals whilst providing constant vascular support. Further research is needed to characterize in detail the function of immune cells within the CNS, describe the metabolic function of vascular cells during chronic hypertension, and investigate systemic inflammatory markers that could predict the onset and progression of cSVD.

Collectively, our findings define the dynamic nature of microglia in the context of cSVD and highlight the stages of microglial phenotypes in the hippocampus and in the cortex along hypertensive states **(Fig. 8)**. Additionally, these data support the notion that the combination of endothelial alterations during early chronic hypertension, coupled with constant microglial reactivity and the influence of aging, shapes a detrimental synergy that contributes to the development of cSVD. These insights enhance our understanding of microglial behavior and emphasize the demand for region-specific investigations to further discern the complex effects of hypertensive states.

**Fig. 8.**
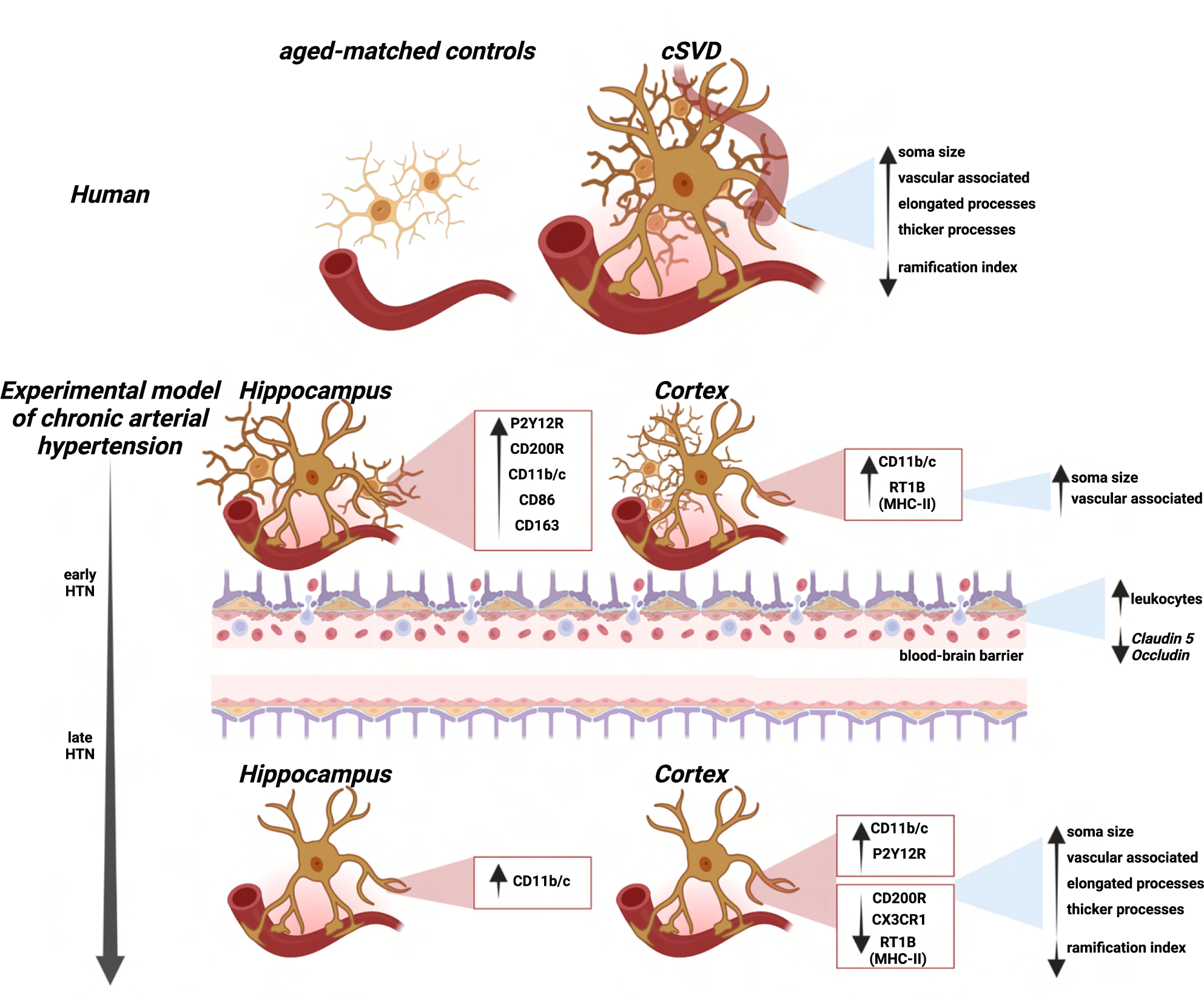
Summarizing figure. Stage-dependent microglia dynamical changes in early and chronic hypertensive states.

### Strengths and limitations

Extensive analysis of microglia in human post-mortem cSVD tissue was conducted, providing valuable insight into distinctive microglial features in the context of cSVD pathology. By utilizing a murine model, we were able to investigate further and validate the observations made, enhancing the reliability and robustness of the results. This approach strengthens the translational relevance of the study. Furthermore, microglia heterogeneity was investigated by integrating high-resolution morphological variables and single-cell surface protein phenotypic characterization. By employing both supervised and unsupervised analyses, the dynamic changes of the microglia landscape in the hippocampus and cortex of normotensive (Wistar) and chronic hypertensive rats (SHRSP) were examined. To the best of our knowledge, this approach, in the context of chronic arterial hypertension and in rats, fills a significant gap in the current literature. Importantly, these results provide evidence of the diversity within microglia sub-populations that confirms a spectrum of microglia phenotypes associated explicitly with chronic arterial hypertension. This study makes an essential step towards characterizing microglia subsets in normotensive-aged brains as well as in chronic hypertensive states. There are several limitations to be considered in this study. First, the human post-mortem tissue data set had a relatively small sample size, which could limit the generalizability of the findings and the statistical power of the analysis. However, it is essential to acknowledge that the availability of post-mortem tissue of patients solely diagnosed with cSVD without comorbid neurodegeneration is scarce, which contributes to the challenge of obtaining a larger sample size in this specific population. Despite the negligible neurodegenerat i ve pathology in the human brain cases included to the study, however, the hippocampus often showed some AD-related changes in the elderly, which is known to cause changes in microglia phenotype, therefore prompting us to limit our investigations to the cortex. To overcome these obstacles encountered in studying human cSVD and to provide additional insights, a rodent model was utilized for further in-depth analyses. Second, while our study successfully identified distinct structural microglial features in chronic hypertensive states, functional assessments were not included in this investigation. The focus was primarily on the phenotypical changes observed in microglia. Further studies should characterize the functional consequences of the identified microglial alterations in the context of cSVD.

---

**Suppl. Table 1.** – List of postmortem human patient samples used in the study.

| **Randomized Case-Code** | **Age** | **Sex** | **Group** | **cSVD -Type** | **NFT Stage (Gallyas)** | **Aβ Stage** | **Aβ Phase** | **PD (LB)** | **Diagnosis** |
| --- | --- | --- | --- | --- | --- | --- | --- | --- | --- |
| Case1 | 54 | f | Control | n/a | I | B | 3 | 0 | Ovarian cancer |
| Case5 | 65 | m | Control | n/a | I | 0 | 0 | 0 | Myocardial infarction |
| Case4 | 71 | f | Control | n/a | 0 | 0 | 0 | 0 | Left heart failure |
| Case8 | 74 | m | Control | n/a | II | 0 | 0 | 0 | COPD, right heart failure, post-tuberculosis |
| Case9 | 74 | f | Control | n/a | II | A | 1 | 0 | Myocardial infarction |
| Case6 | 49 | m | cSVD | WML | I | A | 1 | 1 | Arterial hypertension, cardiac arrhythmia |
| Case 7 | 57 | f | cSVD | WML | 0 | 0 | 0 | 0 | Non-hodgkin lymphoma |
| Case3 | 69 | f | cSVD | WML | I | A | 1 | 0 | Pontine bleeding |
| Case2 | 81 | f | cSVD | WML | I | A | 1 | 0 | Post-ischemic infarct (MCA-I) |

Aβ – amyloid beta; COPD – chronic obstructive pulmonary disease; cSVD – cerebral small vessel disease; LB – Lewy bodies; MCA – middle cerebral artery; n/a – not applicable; NFT – neurofibrillary tangles; PD – Parkinson's disease; WML – white matter lesions

**Suppl. Table 2.**
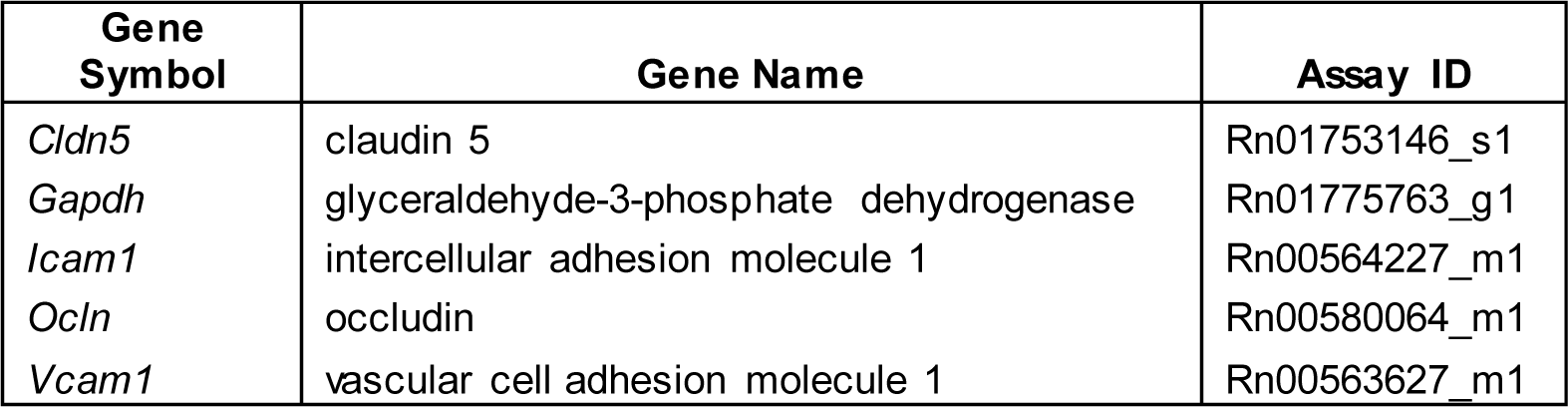
– TaqMan assays used for RT-qPCR analyses.

| **Gene Symbol** | **Gene Name** | **Assay ID** |
| --- | --- | --- |
| *Cldn5* | claudin 5 | Rn01753146_s1 |
| *Gapdh* | glyceraldehyde-3-phosphate dehydrogenase | Rn01775763_g1 |
| *Icam1* | intercellular adhesion molecule 1 | Rn00564227_m1 |
| *Ocln* | occludin | Rn00580064_m1 |
| *Vcam1* | vascular cell adhesion molecule 1 | Rn00563627_m1 |

## Supplementary information

**Suppl. Fig S1.**
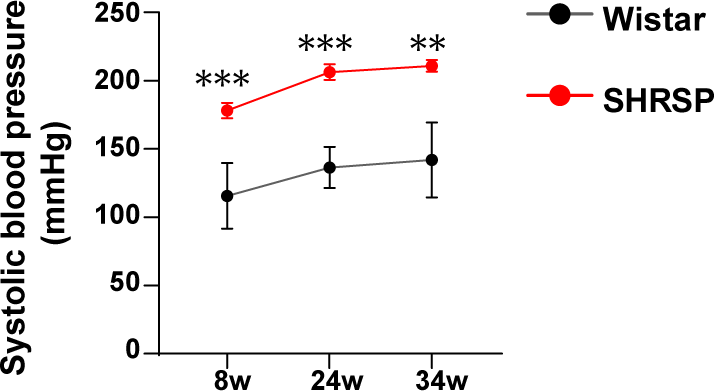
SHRSP exhibit elevated systolic blood pressure from the age of 8-weeks onwards compared to age-matched Wistar controls. Systolic blood pressure was measured in Wistar rats and SHRSP (n = 3 per group) at 8, 24 and 34 weeks of age by indirect tail-cuff method. Pressure and pulse rate signals were continuously recorded and digitalized using BP-2000 Analysis Software (BP-2000 Analysis System, 4-channels, Visitech Systems, Apex, NC, USA). Systolic blood pressure was determined as the mean of ten cuff inflation measurements. Data are represented as mean ± SEM. Statistical analysis was performed using 2way ANOVA with Holm-Sidák’s post hoc test for group and age comparison. *p*-values: ** for *p* ≤ 0.01; *** for *p* ≤ 0.001

**Suppl. Fig S2.**
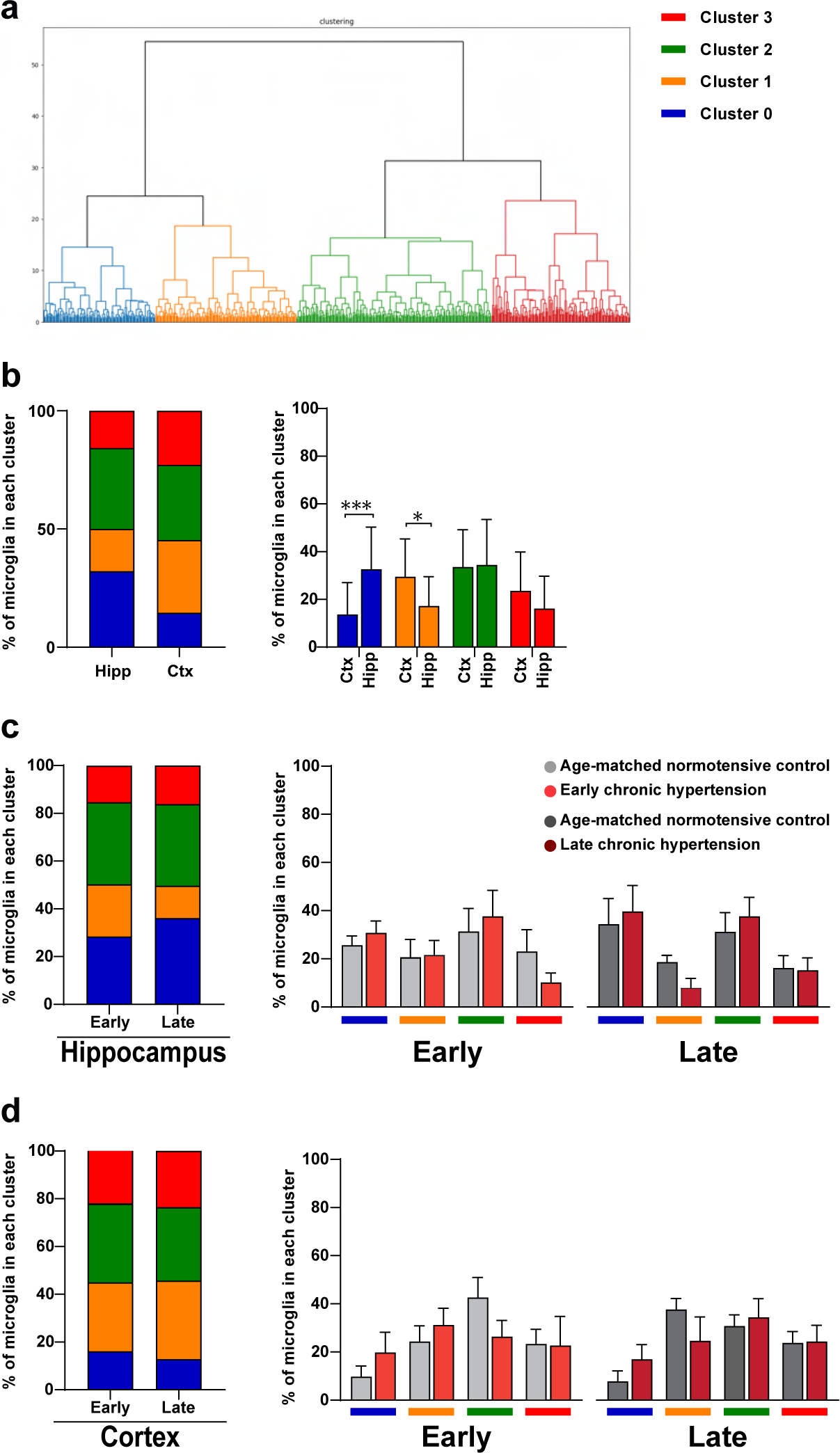
Hippocampal and cortex microglia distribution based on 8 morphological features. **(a)** Ward hierarchical clustering dendrogram of a total of 600 individual Iba1^+^ cells representative of hippocampal CA1 region and retrosplenial cortex captured for individual 3D reconstruction and used in further analysis. **(b)** Relative frequencies of microglia categorized into four distinct morphological clusters in the hippocampus and cortex based on subregions as categorical value**. (c)** Relative frequencies of microglia in the hippocampus and **(d)** the cortex at early and late chronic hypertension. Statistical analysis was performed using 2way ANOVA with Holm-Sidák’s post hoc test for region and cluster comparison. *p*-values: * ≤ 0.05; *** for p ≤ 0.001

**Suppl. Fig S3.**
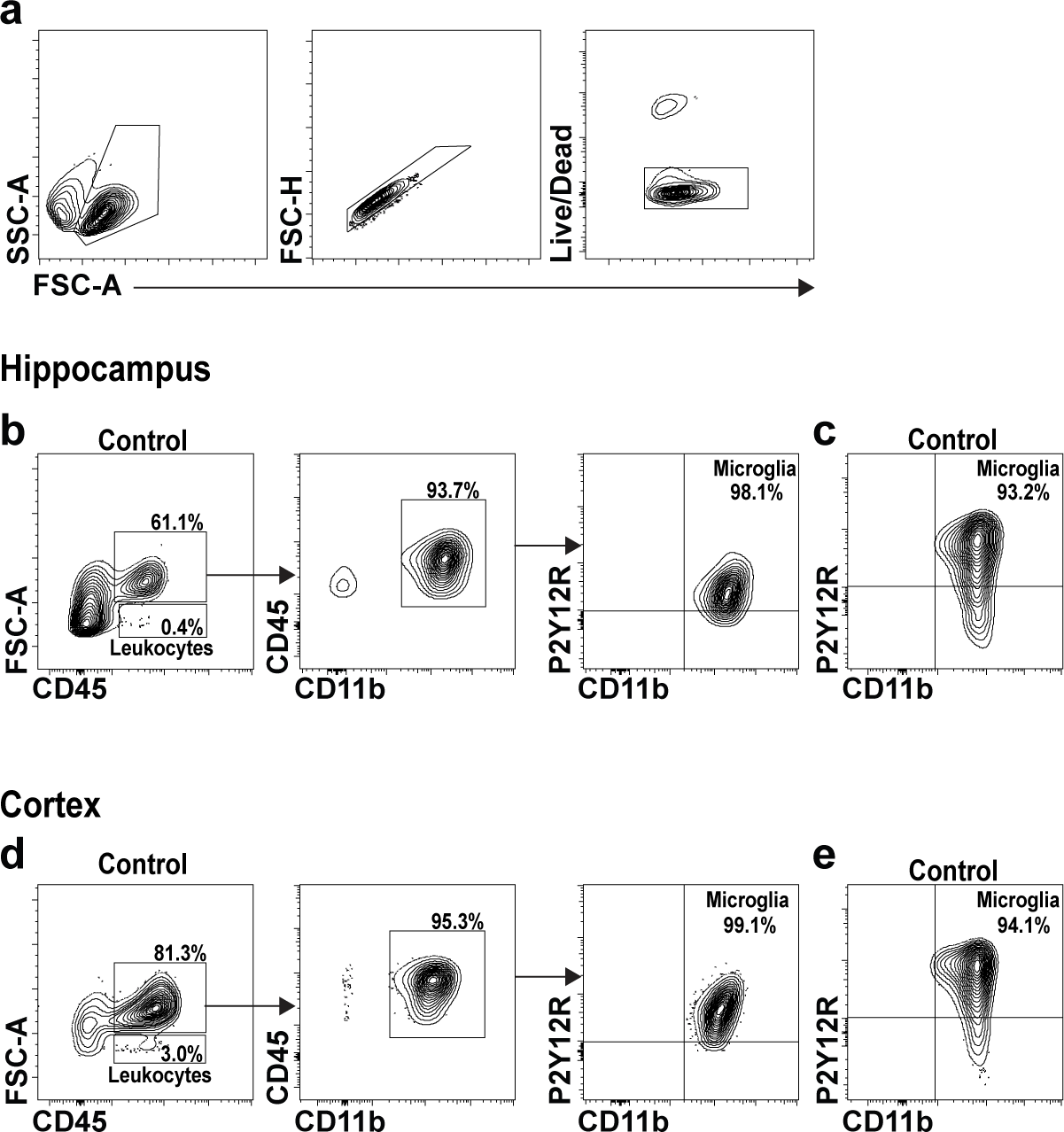
Initial gating strategy and identification of Microglia in normotensive hippocampus and cortex. **(a)** Representative flow cytometric analysis of isolated cells derived from normotensive and hypertensive brains. Cells were selected according to their size and granularity in the forward (FSC-A) and side light scatters (SSC-A). Thereafter, single cells were selected in regard to the ratio of their cell size vs. cell signal displayed in the FSC-H/FSC-A plot. Finally, dead cells were identified by their high affinity to live/dead dye resulting in a brighter fluorescence than live cells. Only live cells were selected for further analysis. **(b, c)** Identification of microglia derived from the hippocampus and (d, e) cortex of normotensive controls via FACS analysis

## Acknowledgements

We thank Petra Grüneberg, Dr. Abidat Schneider, Shaobo Jia (University of Magdeburg), Cornelia Garz (Leibniz Institute for Neurobiology Magdeburg) and Ms. Gabriele Ehmke and Ms. Patricia Häring (Clinical Neuroanatomy, Ulm University) for their expert technical assistance. Figure 8: Stage-dependent microglia dynamics in chronic hypertensive states was created in ©BioRender – Biorender.com.

## Author Contributions

**Conceptualization**: L.M., and I.R.D.; **methodology**: L.M., P.A., A. P. G., S.H., H.M., D. YH., and M.G.; **formal analysis:** L.M., P.A., and D.YH.; **investigation:** L.M., and P.A.; **data curation:** L.M., P.A., and D.YH.; **writing—original draft:** L.M.; **review and editing:** P.A., A. P. G., S.H., A.D., D.YH., S.S., and I.R.D.; **supervision:** S.S., and I.R.D.; **project administration:** L.M., and I.R.D.; **funding acquisition**: S.S., and I.R.D. All authors have read and agreed to the published version of the manuscript.

## Data availability statement

The raw data that supports the conclusions of this article will be made available by the corresponding authors without undue reservation.

